# Decoding of arousal and valence from fMRI data obtained during emotion inductions

**DOI:** 10.64898/2026.03.25.714036

**Authors:** Joel S. White, Yaohui Ding, Nathan M. Muncy, John L. Graner, Leonard Faul, Kevin S. LaBar

## Abstract

Arousal and valence are fundamental dimensions of affective experience signifying levels of activation and pleasantness, respectively. These dimensions play a crucial role in shaping emotional responses and behaviors, with significant implications for psychopathology. Previous machine learning studies had some success decoding these states from brain activation patterns observed during task-based functional magnetic resonance imaging (fMRI), but the results have varied across studies. Moreover, prior studies have often been limited by small sample sizes, weak decoding performance, and non-whole-brain analyses, leaving the neural representations of arousal and valence largely unresolved. Here we successfully decoded arousal and valence from whole-brain task-fMRI data collected from 132 participants during exposure to 300 unique emotional stimuli, including 150 movie clips and 150 text scenarios that reliably induced a wide range of arousal and valence states. Mass univariate general linear models identified block-level activation (emotion stimuli > washout) from all gray matter voxels. Multivariate regression analysis predicted arousal and valence ratings based on these gray matter activations. Patterns in the fMRI data underlying arousal and valence were robust, as they were successfully decoded across both induction modalities using five different linear multivariate regression models. Although significant, decoding from scenarios was less successful than from movies, likely due to their more imaginative nature. In particular, decoding arousal from scenarios only showed low predictive utility. Representations of arousal and valence were widespread throughout the brain, and we reveal cerebellar and brainstem contributions that have largely been absent in past fMRI decoding studies. These findings clarify the distributed neural basis of arousal and valence and provide a foundation for future clinical research on the role of these constructs in affective dysregulation.

## Introduction

### The importance of arousal and valence

The representation and processing of subjective affective states in the brain is a crucial area of study within affective neuroscience. Dimensional theories of emotion propose that all emotional states arise from the combination of two or more basic dimensions of affective processing. For instance, Russell’s circumplex model of affect proposes that emotional states span a two-dimensional space defined by arousal and valence (1). Valence reflects the hedonic quality of an affective state, ranging from unpleasant to pleasant, while arousal reflects the level of activation, ranging from low to high. The circumplex model provides an empirically-supported framework for characterizing a wide range of emotional states along these two neurobiologically and behaviorally distinct dimensions (2). Although these dimensions do not encompass all facets of emotional experience, several theoretical models of emotion emphasize arousal and/or valence as core components of affect (e.g., 3,4).

Abnormalities in the processing of arousal, valence, or both, have been implicated in a variety of neuropsychiatric conditions. Notably, the clinical importance of these constructs is supported by their inclusion as critical domains in psychopathology research in the U.S. National Institute of Mental Health (NIMH) Research Domain Criteria (RDoC) (5). For example, core diagnostic criteria for posttraumatic stress disorder (PTSD) include negative alterations in cognitions and mood, and marked alterations in arousal and reactivity (6). Hyperarousal is implicated in insomnia disorders (7), meltdowns observed in autism spectrum disorder (8), and social anxiety disorders, including social behaviors manifested in Fragile X syndrome (9). Depressive disorders show negative valence biases in cognition in addition to blunted arousal (10), and anxiety disorders are characterized by hypervigilance to threat (11). Collectively, these findings underscore the central role of arousal and valence in shaping emotion-related behaviors across a wide range of neuropsychiatric conditions.

Furthermore, the frequent co-occurrence of mood and anxiety disorders could be explained in the context of shared alterations in arousal and valence. A variety of neuroimaging studies have identified biases towards negative affect within prefrontal and motivation-related circuits that appear across both mood and anxiety disorders (2). These findings suggest that common neural systems may contribute to maladaptive emotional processing across diagnostic categories, further highlighting the importance of characterizing the neural representation of core affective dimensions such as arousal and valence.

### Why multivoxel pattern analysis?

Multivoxel pattern analysis (MVPA) is the application of multivariate techniques, including forms of regression and classification, to fMRI data (12). Multivariate techniques endeavor to build models that can predict a target label (classification) or numerical value (regression) given an associated set of features. Compared to traditional univariate methods, multivariate approaches have higher sensitivity, as they can capture subtle differences in voxel co-activation patterns between task conditions, in addition to the average activation of voxels (13,14). Even if a voxel itself does not change significantly with the target, its signal variability could contribute to a reliable pattern that does. These subtle patterns would be missed by traditional univariate methods. MVPA also facilitates the investigation of affect-related patterns across distributed neural systems at large spatial scales, aligning more closely with modern views of the representation of mental states (15).

### Strengths and limitations of previous MVPA studies

MVPA has previously been used to decode a variety of emotional responses from whole brain fMRI data, including arousal and valence (13). While often successful, interpretation is often limited by small sample sizes, low decoding performance, a within-subject decoding scheme, or a simplified categorical analysis that does not take advantage of the dimensional structure of arousal and valence constructs.

In an early study in this area, Baucom et al. (16) successfully used MVPA to classify fMRI data from 13 participants exposed to affective imagery using four labels: high arousal, positive valence; high arousal, negative valence; low arousal, negative valence; low arousal, positive valence. While successful, the simplified four-way classification limits the interpretation of arousal and valence as dimensional constructs. The classifier was trained and tested within-subject, which can capture participant idiosyncrasies but limits generalization (14). Additionally, the voxel reduction implemented prior to classification could restrict the scope of findings. The small sample size also limits generalizability.

Similarly, Bush et al. (17) used MVPA to classify four groups of high and low arousal and valence from fMRI data collected from 19 participants during exposure to affective imagery. While the study effectively explored inter-study correlations in affective classification, conclusions regarding the neural correlates of arousal and valence were again limited by the simplified four-way classification approach, within-subject scheme, and small sample size. Expanding upon this work, Wilson et al. (18) incorporated concurrent heart rate deceleration into a multivariate regression analysis and successfully decoded valence, demonstrating the potential benefits of integrating physiological signals into affective decoding. However, while statistically significant, the analysis could not explain more than 6% of the total variance of the full valence scores. Performance improved after the exclusion of samples with neutral valence, but this regression on only extremely pleasant or unpleasant stimuli does not represent the full range of the valence dimension.

Kim et al. (19) employed a searchlight analysis to predict valence from fMRI data collected from 17 participants during the viewing of naturalistic stimuli, specifically a 48-minute TV episode. The study divided valence into signed (positive-to-negative) and unsigned (valenced-to-non-valenced) components, and successfully extended previous results to naturalistic settings. However, while statistically significant, the analysis only weakly predicted signed valence and could not predict unsigned valence, potentially due to an oversaturation of neutral arousal and valence values and a limited number of extremes. Additionally, the small sample size (N=17) and exclusion of the cerebellum and brainstem limit interpretability and generalization of the results.

Kim et al. (20) successfully used a deep neural network (DNN) to decode both arousal and valence from fMRI data collected from 10 male participants during exposure to affective sounds. While some research has suggested that non-linear models, such as a DNN, perform similar to, if not worse than, linear models with fMRI data (21), non-linear models can potentially identify higher-order relationships within the dataset (15). The high performance achieved with the DNN using a within-subject scheme revealed regions associated with the processing of arousal and valence and demonstrated the utility of DNNs. However, the interpretation of these neural correlates may be limited by the small, all-male sample size and the complexity of the DNN. In addition to a DNN, non-linear and linear support vector machines were implemented, although neither of which produced significant decoding performance for either arousal or valence.

Although this review is not exhaustive, past arousal and valence decoding studies were often limited by small sample sizes, an analysis which simplifies arousal and valence from continuous dimensions to coarse categories, a feature reduction approach which under-represents areas like the cerebellum, a within-subject training scheme which limits generalizability, and weak or insignificant decoding performance. Moreover, previous studies often only reported results from one type of multivariate model, raising questions regarding how robust the captured fMRI patterns were to variation in modeling parameters. A recent 2025 meta-analysis found that no valence decoding fMRI study except Kim et al. (20) evaluated multiple machine learning models on the same dataset, although different models impose distinct feature structures and can differ substantially in their ability to capture predictive signals and in the neural patterns they detect (22). Finally, previous studies have relied on one modality to induce emotional states, leaving open the question of how representations of arousal and valence generalize across types of stimuli.

These prior limitations motivated the design of the present study. We collected fMRI data from a large sample size with two induction modalities (movies: final N=114, 66 female; scenarios: final N=121, 67 female). We implemented multiple multivariate models, and used a regression approach to treat arousal and valence as continuous dimensions. Each MVPA was conducted with a group-level decoding scheme to improve generalizability. Lastly, gray matter voxels from the entire brain, including the cerebrum, cerebellum, and brainstem, were used as MVPA features, ensuring that all possible informative voxels can contribute to decoding.

We hypothesized that we would achieve significant decoding performance across a range of multivariate models. Furthermore, we hypothesized that the importance maps would be widely distributed throughout the brain, including cerebellar, brainstem, and other subcortical regions often undersampled in prior work. The cerebellum, in particular, has rarely been reported in prior fMRI arousal and valence decoding studies, but a growing body of multimodal evidence has suggested that numerous regions of the cerebellum are involved in a wide array of emotional and higher-order cognitive processes (23,24). The sensitivity of MVPA to subtle neural patterns, combined with our large sample size and smaller voxel size relative to prior work, allowed us to more closely evaluate the role of more elusive contributions of smaller brain regions to arousal and valence. We additionally expected modest overlap between induction modalities in heteromodal association cortices and a significant contribution of ventromedial prefrontal cortex (vmPFC) to valence processing, given its established role in prior work across both univariate and multivariate methods (e.g., 25,26). While individual regions like the vmPFC may show up across studies, multivariate methods leverage the contribution of each voxel in an interconnected fashion; therefore findings are better interpreted in the context of distributed patterns rather than isolated regional effects. Prior decoding studies have implicated a number of regions in arousal and valence processing, though consensus varies. Building from regions reported more consistently, we expected our valence MVPA to utilize the ventrolateral prefrontal cortex (vlPFC) and anterior cingulate cortex (ACC), and our arousal MVPA to utilize dorsolateral prefrontal cortex (dlPFC), as suggested by prior decoding studies (16,20).

## Results

### Arousal and valence target ratings

The arousal and valence block-level ratings act as the targets in the MVPA. Fig 1 depicts the arousal and valence block ratings by modality. Each point on the graph represents the group-averaged ratings for a single block of trials. Block ratings do not fully separate into the original 15 emotions in a space of arousal and valence, although notable clusters can be observed. For example, in the movie ratings, some overlap appears between fear and anxiety, and between amusement, awe, craving and romance.

**Fig 1.**
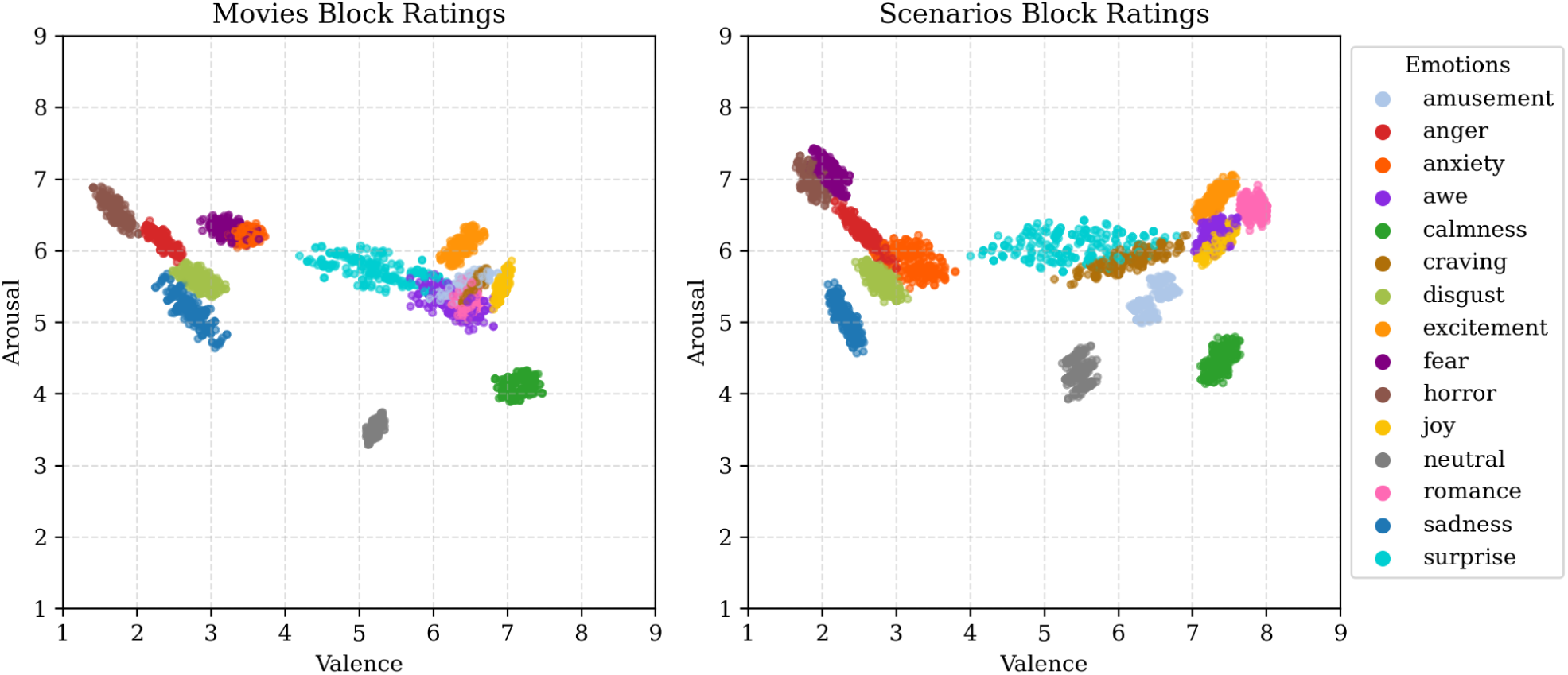
Scatter plots depicting all arousal and valence block ratings for movies (left) and scenarios (right). Points are colored according to their original categorization of 15 emotions, which are shown in the legend on the right, although these categorical ratings were not utilized in the present analyses.

### Successful decoding of arousal and valence

As depicted in Fig 2, rated and predicted arousal and valence ratings from movie decoding test splits are significantly correlated in all five implemented models, as calculated by permutation testing. The *p*-values and mean *R*^2^ and Pearson’s correlation coefficients for each of 25 model repetitions, with standard deviations calculated across repetitions, are as follows: for arousal, *p* < 0.001; *r* = 0.558, *SD* = 0.003; *R*^2^ = 0.301, *SD* = 0.004 and for valence, *p* < 0.001; *r* = 0.582, *SD* = 0.004; *R*^2^ = 0.330, *SD* = 0.005, as achieved with the Ridge model. Prediction patterns between models are relatively similar but differ slightly. The models can overestimate lower values and underestimate higher values, reflecting the error-penalization strategy used during model tuning to achieve optimal overall performance.

**Fig 2.**
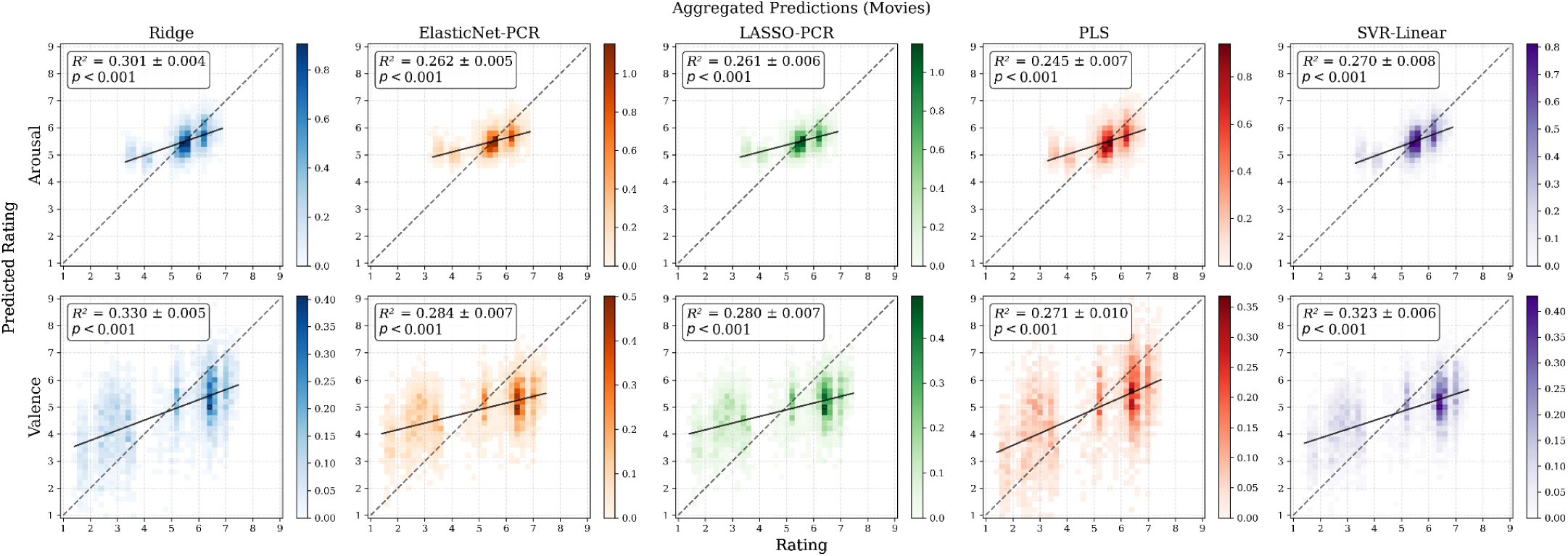
2D histograms of the predicted versus rated arousal and valence values for each movie prediction model across all five test sets of a sample five-fold cross-validation. The included *R*^2^ values were averaged across 25 repetitions of each model. The *p*-values were estimated empirically against 1000 repetitions of permutation tests, making a *p*-value of approximately 0.001 the smallest achievable. The x-axis represents the rated score, and the y-axis represents the model’s predicted value. Colorbars are defined using density, which represents the ratio of values present in a bin, divided by the bin size, which is 0.25 in this case. The dotted line represents a perfect one-to-one correlation.

The decoding of arousal and valence from scenarios was less successful than movies, but still statistically significant for both arousal (p < 0.001; r = 0.156, *SD* = 0.006; *R*^2^ = 0.023, *SD* = 0.002) and valence (p < 0.001; r = 0.381, *SD* = 0.005; *R*^2^ = 0.141, *SD* = 0.004), as achieved with the Ridge model. 2D histograms of the predicted versus rated arousal and valence values for each model when decoding scenarios can be found in Supporting Information, “Example scenario decoding predictions.” As shown in Fig 2 and Fig 3, success was replicated across all five models in the decoding of both arousal and valence from fMRI data collected during movies. Ridge outperformed all other models in both arousal and valence decoding for both movies and scenarios, as shown in Fig 3; see Supporting Information, “Model performance versus permutations” for comparisons of model performance to their associated null permutations.

**Fig 3.**
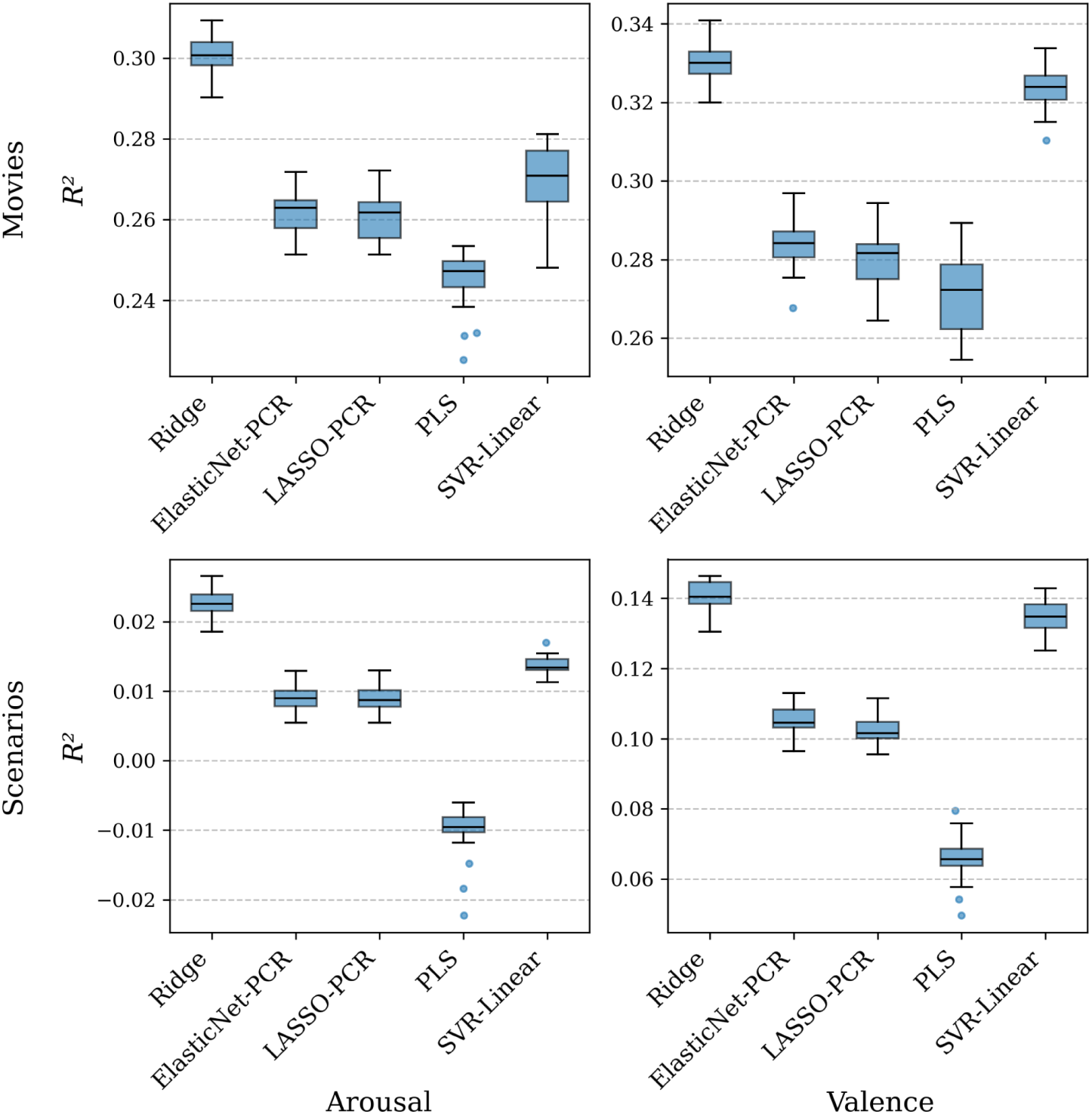
Boxplots depicting the average *R*^2^ values across the 25 repetitions of each model. The top row shows model performance in movie decoding and the bottom row in scenario decoding. The left column shows arousal decoding, and the right column shows valence decoding. All models were significant by permutation testing with 1000 repetitions, with all achieving a *p*-value less than 0.001, except decoding of arousal with scenarios by PLS, which achieved a *p*-value of 0.026. Performance relative to permutations can be found in Supporting Information, “Model performance versus permutations.”

In decoding arousal with data collected during scenario viewing, several models produced statistically significant permutation tests despite yielding cross-validated *R*^2^ values close to zero or even negative. A negative *R*^2^ implies the model’s summed squared prediction error exceeds the error of a constant predictor that always returns the training-set mean; in other words, the model performs worse than simply guessing the mean for every block. Permutation significance in this case indicates the model’s predictions are non-randomly related to the labels, but the relationship is too weak or noisy to provide useful predictive accuracy under the *R*^2^ metric. Because statistical significance, as in permutation *p*-value, and predictive utility, as in *R*^2^, convey different information, we report both and caution that a significant *p*-value alone does not imply a useful model. Therefore, while significant, the low *R*^2^ values indicate that the decoding of arousal from scenarios was weak; as such, the corresponding neural correlates are presented in Supporting Information, “Full Ridge visualizations” rather than the section below.

### Distributed neural representations of arousal and valence

While regions are reported individually for clarity, these results reflect multivariate patterns distributed across the brain. MVPA decoding utilizes the combined, conditionally-dependent contribution of each region in an interconnected fashion rather than the independent effect of any single region. A positive weight on a voxel means that, in the context of the full MVPA, that voxel pushes the prediction towards the positive end of the target dimension. The sign of the weight does not necessarily imply a direct increase or decrease in activation for a block, as a univariate analysis would. While not the main goal of this study, a univariate analysis of the data can be found in Supporting Information, “Parametric modulation analysis,” and showed patterns generally similar to that of the MVPA. Important regions from the Ridge model are reported here, as it achieved the highest decoding performance across both arousal and valence in both tasks, and full slices and MNI coordinates of all significant clusters for the Ridge model can be found in Supporting Information, “Full Ridge visualizations”. While all models rely on multivariate patterns, the specific MVPA algorithm determines how that inter-regional structure is captured, leading to subtle differences in the spatial patterns of significant voxels. Therefore, full visualizations and cluster tables for all five implemented models can be found in the code repository described in the Reproducibility section.

As depicted in Fig 4, voxels important in the decoding of arousal and valence from fMRI data collected during movie viewing demonstrated distributed neural patterns related to the prediction of emotional responses. Regions important to the decoding of arousal from fMRI data collected during movies included a number of cortical, subcortical, cerebellar, and brainstem regions. Cortical clusters were observed in the dorsolateral prefrontal cortex (dlPFC) and the posterior cingulate cortex (PCC), among regions broadly associated with sensory processing, including the precuneus and the primary visual area. In the subcortex, significant voxels appeared in the mesencephalic reticular formation (MRF) and the thalamus. Lastly, in the cerebellum, important voxels for decoding appeared in multiple regions along the vermis, including vermis IX, vermis crus II, and vermis VI, in addition to medial VIIb and lateral crus I/II. All clusters appear bilaterally, if not along the midline.

**Fig 4.**
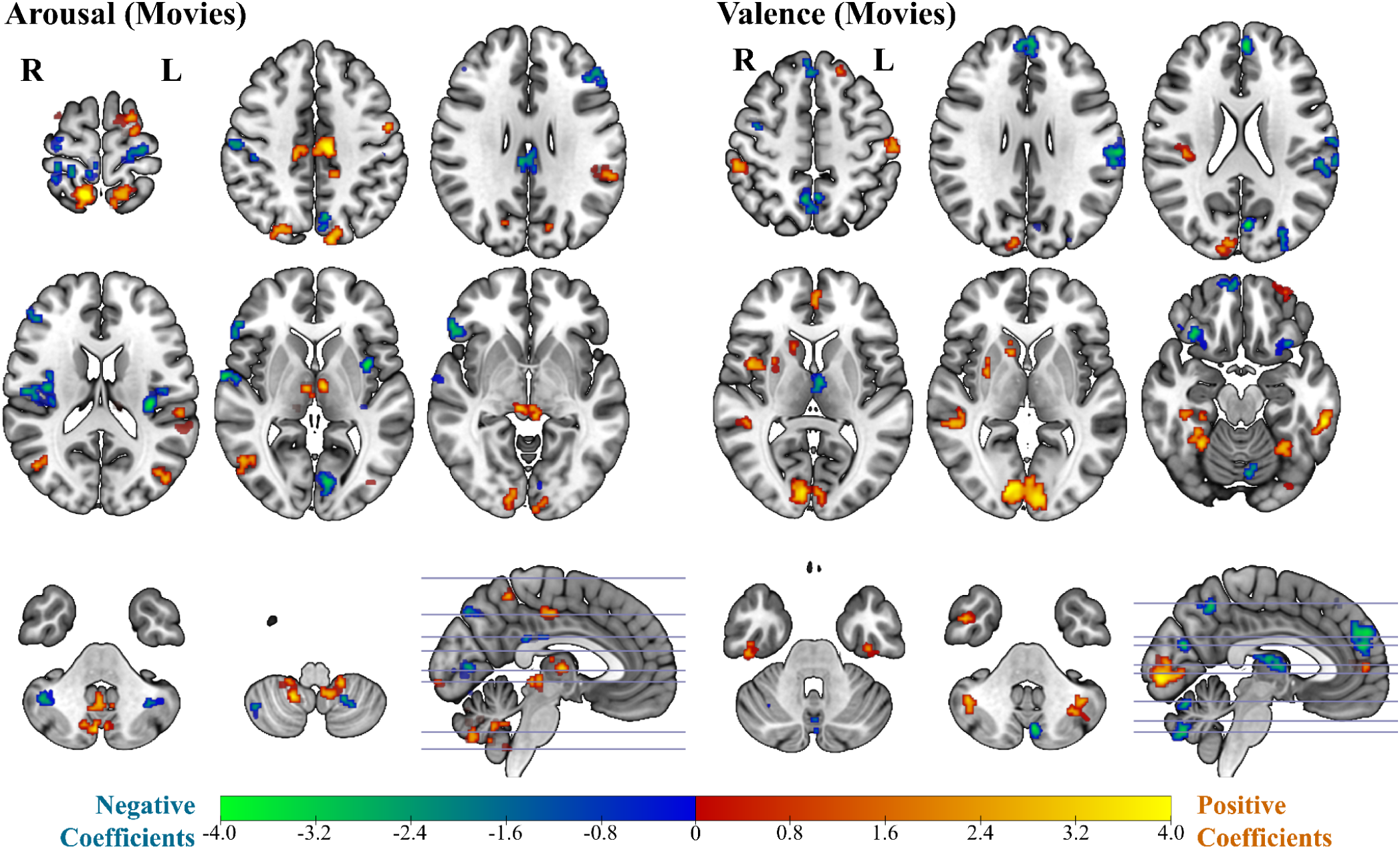
Slices depicting important voxels from the Ridge regression model used to decode arousal (left) and valence (right) from fMRI data acquired during movie viewing. Voxel-wise model coefficients are thresholded at |z| > 1.96, and only clusters of ≥30 contiguous voxels (NN=1) are shown. Negative coefficients are displayed in blue-green, positive in red-yellow. Visualization was created with MRIcroGL (27). A full set of evenly-spaced slices and MNI coordinates of all significant clusters for the Ridge model can be found in Supporting Information, “Full Ridge visualizations.”

The decoding of valence from fMRI data collected during movies also spanned cortical, subcortical, and cerebellar regions. Cortical clusters included the dorsomedial prefrontal cortex (dmPFC), ventrolateral prefrontal cortex (vlPFC) extending into the anterior insula, right medial orbitofrontal cortex (mOFC), paracingulate gyrus, and right parahippocampus (PHC). Additionally, the left thalamus, VI, and crus I/II were important for decoding.

Voxels important in the decoding of valence from fMRI data collected during scenario viewing display distinct neural patterns related to the prediction of emotional responses, as shown in Fig 5. The Ridge model’s significant voxels primarily included regions within the cortex and cerebellum. Cortical clusters included the dmPFC, the ventromedial prefrontal cortex (vmPFC), multiple sections of the anterior cingulate cortex (including the pregenual anterior cingulate cortex (pgACC), subgenual anterior cingulate cortex (sgACC), and dorsal anterior cingulate cortex (dACC)), right vlPFC, left posterior mOFC, and right PHC. Significant cerebellar voxels were observed in crus I/II, including individual clusters within the intermediate and lateral portions.

**Fig 5.**
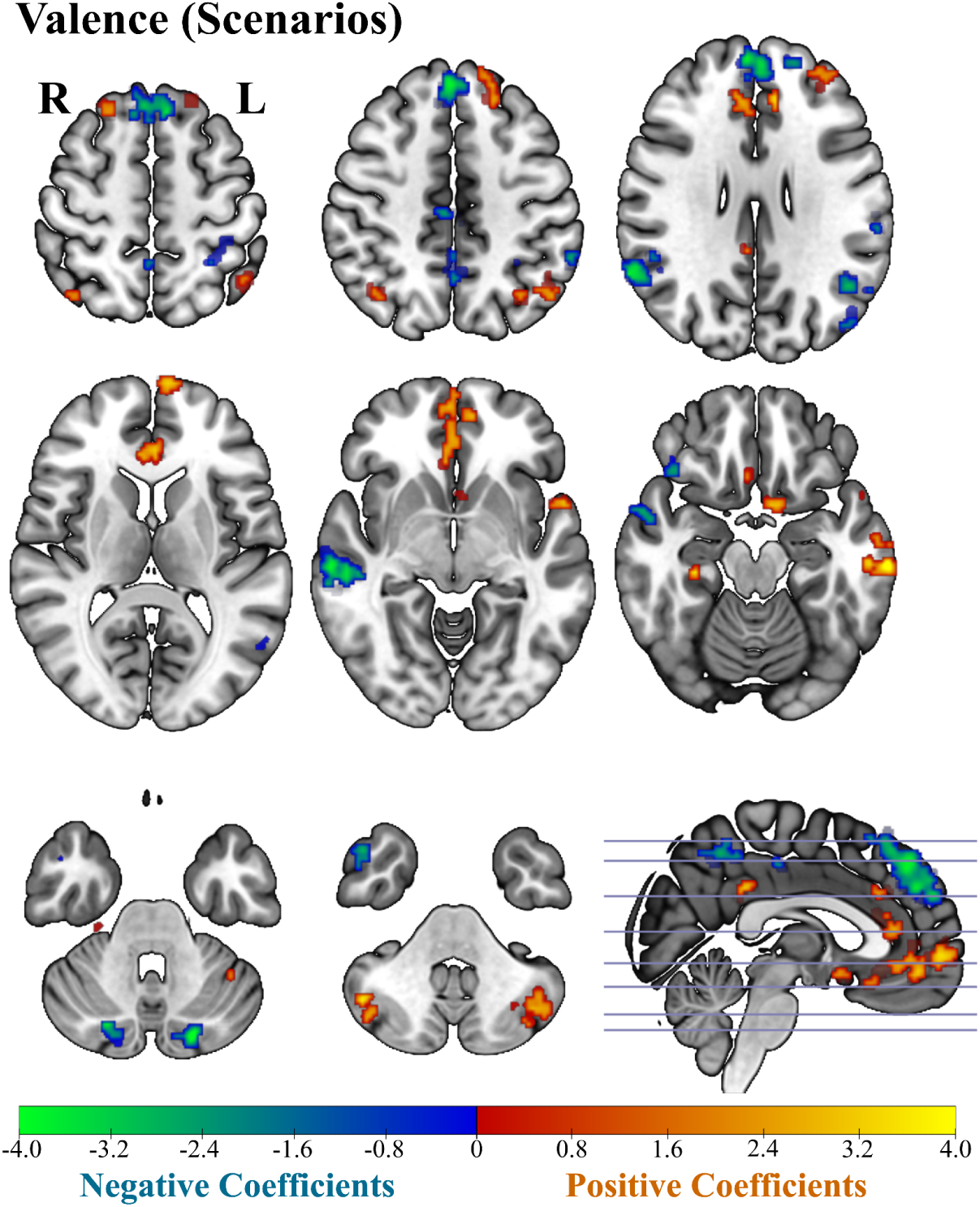
Axial slices depicting important voxels from the Ridge regression model used to decode valence from fMRI data acquired during scenario viewing. Voxel-wise model coefficients are thresholded at |z| > 1.96, and only clusters of ≥30 contiguous voxels (NN=1) are shown. Negative coefficients are displayed in blue-green, positive in red-yellow. Visualization was created with MRIcroGL (27). Full slices and MNI coordinates of all significant clusters for the Ridge model can be found in Supporting Information, “Full Ridge visualizations.”

Comparing important regions from the decoding of valence between movies and scenarios sessions revealed both substantial overlap and task-specific patterns. Shared regions included the dmPFC, vlPFC (although vlPFC in the scenarios results was restricted to the right hemisphere), mOFC (present in both but restricted to the left posterior mOFC in scenarios), right PHC, and cerebellar crus I/II. Scenario-specific clusters included the vmPFC in addition to multiple ACC subregions, specifically the pregenual, subgenual, and dorsal ACC. The movie data uniquely contains a cluster in the left thalamus. Additionally, voxel importance differs in sensory areas, with movie decoding utilizing a large cluster in V1/V2 for example.

## Discussion

Arousal and valence are foundational dimensions of human affect with broad implications towards affect processing in both healthy and clinical populations. Prior decoding studies have been limited by small sample sizes, non-whole-brain analyses, binarization of continuous affective ratings, within-subject training schemes that limit group inference, weak or inconsistent decoding performance, reliance on a single MVPA model, and/or use of a single stimulus modality. To address these limitations, we decoded whole-brain fMRI data from 132 healthy adults during emotion inductions using either short movie clips or text scenarios. We trained and evaluated multivariate regression models at the group-level using continuous arousal and valence ratings. Across multiple modalities and multivariate models, we successfully decoded both arousal and valence. Although regression models were significant in all cases, the variance explained by models decoding arousal from scenarios was negligible. By examining which regions the models assigned the greatest importances, we identified neural representations of arousal and valence spanning cortical, subcortical, and cerebellar regions.

### Replicated and extended neural representations

Regions implicated in the representations of arousal and valence are somewhat consistent with previous decoding studies, but additionally extend the literature, particularly by implicating additional subcortical and cerebellar regions. It should be kept in mind that MVPA utilizes distributed neural patterns, rather than the independent effects of individual regions and so results should always be interpreted within the context of the broader set of voxel importance maps.

Consistent with prior arousal decoding studies, our decoding of arousal from movies implicated the dlPFC (16), additionally aligning broadly with prior reports of middle frontal gyrus involvement (20). The PCC was also identified, which was associated with arousal in a prior decoding study and implicates default mode network involvement (20). Our findings reveal a role for other brain regions that expand upon those identified in prior arousal decoding studies and may extend the current understanding of the neural basis of arousal. In particular, the MVPA analysis identified the MRF, a brainstem region associated with regulating attentional vigilance and threat perception (28), and the thalamus, which similarly modulates attention in addition to integrating and relaying emotional information (29). Additionally, prior decoding studies have underrepresented the cerebellum and its integration with affect and cognition. Numerous cerebellar clusters were identified by the MVPA analysis, and are supported by the literature. For instance, Styliadis et al. (30) examined the processing of arousal and valence in the cerebellum using magnetoencephalography (MEG) and found arousal to be processed within vermis VI and left medial crus II, matching the vermis VI and bilateral crus I/II clusters identified in our analysis. Our analysis also implicated vermis IX, often referred to as the “limbic cerebellum” given its involvement in affective regulation (23,24,31). Lastly, the involvement of crus I/II aligns with prior research linking these regions to emotional-cognitive integration (23,32).

The decoding of valence from movies identified the vlPFC, consistent with prior valence decoding studies (16,20). This cluster in the vlPFC additionally extends into the anterior insula, a region revealed in a prior decoding study (20), and a meta-analysis examining the brain basis of positive and negative affect (26). The MVPA analysis selected voxels in the right mOFC, previously identified via valence decoding with a deep neural network by Kim et al. (20), and right PHC, previously identified with MVPA classification by Baucom et al. (16). Our MVPA additionally identified the dmPFC, a region associated with social and emotional processing; the model’s high-magnitude negative coefficients in the dmPFC imply a contribution towards negative valence, which aligns with this region’s inclusion in the multivariate Picture Induced Negative Emotion Signature (33). The model also included a cluster in the left anterior medial thalamus, which is part of a hippocampal-thalamo-PFC circuit important for emotional, mnemonic, and executive functions (34). The cluster in the paracingulate gyrus may contribute to positively-valent emotional processing through self-referential processing. Lastly, the model showed bilateral clusters in crus I/II and superior VI. This pattern is consistent with Styliadis et al. (30), who used MEG to show that low valence was processed in left VI, and, when high valence interacted with high arousal, left crus I.

The decoding of valence from scenarios overlapped to a large extent with that of movies, but the difference between modalities did manifest in notable differences within sensory and prefrontal areas. It should be noted that while valence was successfully decoded from both modalities, the model that decoded valence from movies showed higher predictive utility than that of scenarios. We believe this modality difference in decoding performance can be attributed to the imaginative nature of scenarios, which entirely relies on the participants’ generative ability to imagine themselves in a textual scenario, as opposed to movies, which guides participants with visual input. A scenario-specific cluster appeared in the vmPFC, which could be involved in subjective evaluation, and therefore appear as participants attempt to mentally create a valenced state using the text scenario. This cluster in the vmPFC, together with the cluster in the dmPFC, mirrors prior evidence linking the vmPFC to appetitive value and the dmPFC to aversive value processing in a review of valence and decision making (35). Furthermore, scenario-specific clusters appeared in the pgACC, sgACC, and dorsal ACC, regions which may integrate emotional appraisal, self-referential processing, and the regulation of positive and negative affect (36). Movie decoding contains significantly more sensory-related voxels than scenarios, including, for example, a large cluster spanning the primary and secondary visual cortices. Movie decoding also contains a cluster in the left thalamus. Both movies and scenarios contained clusters in the dmPFC, vlPFC, mOFC, right parahippocampus, and cerebellar crus I/II, which together may contribute to valenced states independent of modality.

Notably, all of the top performing multivariate models contained significant clusters in the cerebellum, a structure largely underrepresented in previous arousal and valence decoding studies. More specifically, the MVPA selected the crus I/II to aid in decoding both arousal and valence with fMRI, with bilateral clusters appearing in Ridge decoding of arousal and valence from movies, in addition to valence from scenarios, suggesting a broad and potentially modality-independent role in affective processing. Crus I/II in particular have been found to project to distributed prefrontal and limbic regions (23,32). These findings add to a growing body of multimodal evidence suggesting the cerebellum supports higher-order emotional and cognitive functions beyond its traditional role in motor control (23,24,31,37).

### Successful decoding with multiple linear MVPA models

Some previous studies have reported low, but significant, performance decoding arousal or valence using linear MVPA models. Kim et al. (20) found insignificant performance using Linear-SVM to decode arousal and valence from whole-brain fMRI data, whereas a deep neural network succeeded. The study suggested that extracting meaningful features from whole-brain fMRI data would be difficult without a multiple-layer architecture such as a deep neural network (20), and noted that classifying multivoxel patterns across the whole-brain with a single-layer model such as SVM often involves feature extraction, which has been employed in a number of prior decoding studies (16,22,38). In contrast, the present study found that several different linear models could all successfully decode both arousal and valence from movies, and with correlation and *R*^2^ values exceeding those of all examined prior studies. However, while statistically significant, decoding from scenarios was less effective, and the predictive utility of arousal, in particular, was negligible. We believe this difference in decoding performance stems from the imaginative nature of scenarios. While movies guide participants with visual input, the effectiveness of scenarios relies entirely on participants’ ability to imagine themselves within the described situation. Additionally, successful decoding was achieved with input from all gray matter voxels in the brain, rather than only pre-selected regions. However, it should be acknowledged that providing all gray matter voxels, 158,110 in this case, as features to the MVPA substantially increased computational complexity. See Supporting Information, “MVPA computation efficiency,” for notes regarding computational efficiency and the hardware required to run these models.

## Limitations and future directions

The present study implemented five different linear multivariate models. However, non-linear models could, in principle, capture more complicated relationships within fMRI data, although at the expense of computational efficiency, which already suffers due to the high dimensionality of neuroimaging data, and interpretability. Additionally, the target arousal and valence values were derived from post-scan ratings obtained during a second exposure to the stimuli. Because affective responses can shift across repeated viewings, these ratings may not have perfectly captured participants’ initial in-scanner affective response.

Future work could test higher-dimensional models of emotion beyond the arousal-valence framework, for example by adding a dominance or motivation dimension. Future work could also explore additional modalities of emotion induction, such as music or naturalistic stimuli, and investigate how neural representations differ across contexts. The present study identified shared regions representing valence across movie and scenario inductions, but the interpretation of this is made difficult by differences in decoding performance, with movies being decoded better than scenarios, especially for arousal.

This study was conducted in adults between the ages of 18 and 49. Therefore, the generalizability of the identified neural representations of arousal and valence to children or older adults remains uncertain. Similarly, the application of identified neural representations of arousal and valence could be extended to clinical populations, as supported by their inclusion as critical domains in psychopathology research within the NIMH RDoC initiative (5). Considering that abnormalities in the experience of arousal and valence underlie numerous psychiatric disorders, an understanding of the brain mechanisms supporting these constructs is therefore paramount to characterizing affective psychopathology across a broad spectrum.

## Conclusions

We successfully decoded both arousal and valence from fMRI data in a large sample size of healthy adult participants (N = 132) during emotion induction using both short movie clips and one-to-two sentence text scenarios. Patterns in the fMRI data were robust, and successfully decoded by five different multivariate regression models: Ridge, ElasticNet-PCR, LASSO-PCR, PLS, and Linear-SVR. The highest performing model, Ridge, achieved Pearson’s correlation coefficients of (mean ± standard deviation) 0.558 ± 0.003 when decoding arousal from movies, 0.582 ± 0.004 for valence from movies, 0.156 ± 0.006 for arousal from scenarios, and 0.381 ± 0.005 for valence from scenarios, all of which were significant by permutation testing at *p* < 0.001. Decoding from scenarios was slightly less effective compared to movies, likely due to their imaginative nature. By examining which regions the model assigned the greatest importances, we identified the neural representations of both arousal and valence, and, in the case of valence, across modalities (since arousal explained little variance in response to scenarios). The neural representations of arousal and valence consisted of clusters broadly distributed across the cerebrum, many of which replicated findings of previous studies, in addition to numerous prominent clusters in the cerebellum and brainstem, which align with the broader literature but have largely been absent in many prior decoding studies. These neural systems of arousal and valence provide a foundation with which future research can investigate altered affective processing, including that which underlies symptomatology across various psychopathologies.

## Materials and methods

### Participants

A total of 1257 individuals were recruited from the local Durham, NC population. 166 participants were enrolled in the study. An additional 5 pilot participants completed sessions before the task was finalized and are not included in analyses. A total of 132 healthy English-speaking adult participants completed at least one session of the study with fMRI data that passed quality control (including checks for excessive motion and other artifacts). The participants were between 18 and 49 years old (*N* = 132; 56.1% female; *M*_age_ = 30.05 years; *SD*_age_ = 8.39 years; 6.1% left-handed). Recruitment was designed to achieve a sample broadly representative of the demographics of the local Durham, North Carolina, USA, area. Participants were required to have completed at least a high school diploma or equivalent qualification. Exclusion criteria included: a history of neurological conditions, current psychiatric disorder diagnosis, current use of psychotropic medications, current substance use disorder, cardiovascular or respiratory illness, or contraindications to research MRI (intracranial or other medical implants, transdermal drug delivery systems, non-removable body piercings, pregnancy, or claustrophobia). Each participant provided written informed consent in accordance with the Duke University Institutional Review Board ethical standards (Protocol Number: Pro00105943). Compensation was provided at a rate equivalent to $25/hour. Fig 6 details information regarding excluded participants and data by session along with participant inclusion by modality. After applying fMRI quality control criteria, 114 participants had data for the movies session of sufficient quality for analysis (57.9% female; *M*_age_ = 29.98; *SD*_age_ = 8.42) and 121 participants had usable data for the scenarios session (55.4% female; *M*_age_ = 30.04; *SD*_age_ = 8.49). Across these participants, 3143 blocks of movies and 3247 blocks of scenarios were retained for multivariate analysis, whereas 277 movie blocks and 383 scenario blocks were excluded for excessive motion.

**Fig 6.**
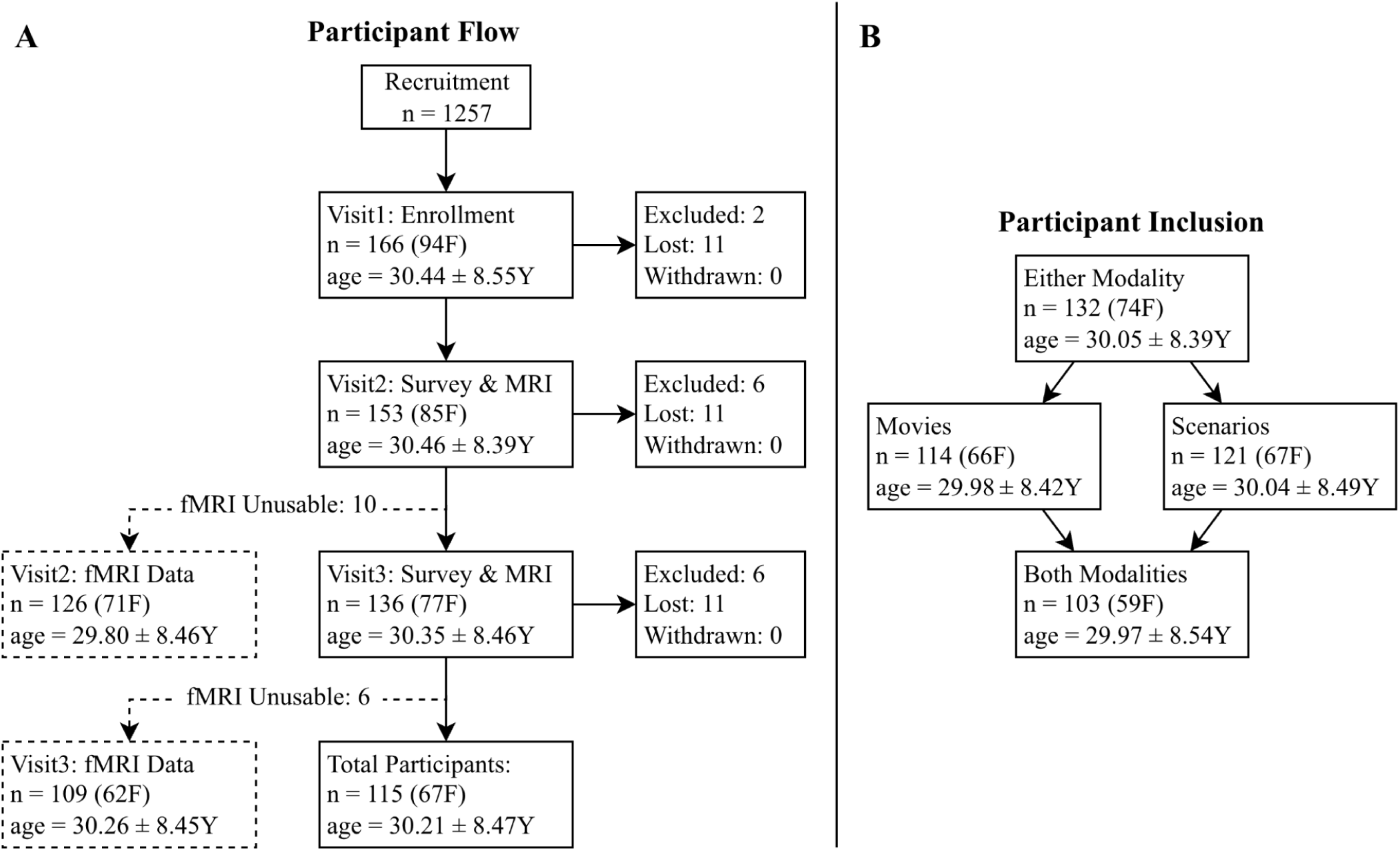
Participant recruitment flowchart (left) and participant inclusion diagram (right). (A) Participant recruitment and retention flowchart by session. A total of 1257 individuals were screened for eligibility. Of those, 166 participants were enrolled in the study (56.6% female (F); *M*_age_ = 30.44 years (Y); *SD*_age_ = 8.55). A total of 115 participants completed all sessions of the study (58.3% F; *M*_age_ = 30.21 Y; *SD*_age_ = 8.47). All but 10 participants’ second visits and all but 6 participants’ third visits had fMRI data that passed quality control. Considering participants had varying ordering of movie and scenario induction sessions, this flowchart only reflects retention over time as opposed to inclusion by modality. (B) Participant inclusion by induction modality. After applying fMRI quality control criteria, 114 participants had data for the movies session of sufficient quality for analysis (57.9% female; *M*_age_ = 29.98; *SD*_age_ = 8.42) and 121 participants had usable data for the scenarios session (55.4% female; *M*_age_ = 30.04; *SD*_age_ = 8.49).

### Experimental design

#### Emotion induction task

Participants performed an emotion induction task while undergoing functional MR imaging in a 3.0T Siemens Magnetom scanner. Participants completed the task in two sessions (separated by at least seven days), viewing emotional video clips during one of the sessions and reading emotional text scenarios during the other session (order was counterbalanced across participants). The stimuli were composed of 150 externally-validated, emotionally-evocative, short (3 to 8 sec duration), mute movie clips (39) and 150 externally-validated, two-sentence, second-person text scenarios depicting hypothetical emotional situations (40,41). Stimuli from both modalities were divided into 15 emotion categories, based on normative ratings, with 10 stimuli per category per modality. Stimuli were never repeated. The set of emotions included amusement, anger, anxiety, awe, calmness, craving, disgust, excitement, fear, horror, joy, neutral, romance, sadness, and surprise. See Supporting Information, “Movie and scenario stimuli” for a more detailed description of the stimuli. A single trial consisted of viewing a movie or scenario and mentally replaying the stimulus once it disappeared from the screen, which, combined, lasted 13 seconds, followed by a location judgement question about where the contents of the stimulus took place (indoor/outdoor). In each session, participants experienced 30 blocks, each consisting of 5 trials with stimuli from the same emotion category. At the end of each block, participants were asked to endorse which emotion they experienced the most during that block and then to what degree they experienced that emotion, on a scale from 1 to 10. A neutral washout image separated the blocks. Following each scanning session, participants rated the arousal and valence of half the stimuli shown during that session. The task is depicted in Fig 7. Further details about the task and recording hardware can be found in Supporting Information, “Full task description.”

**Fig 7.**
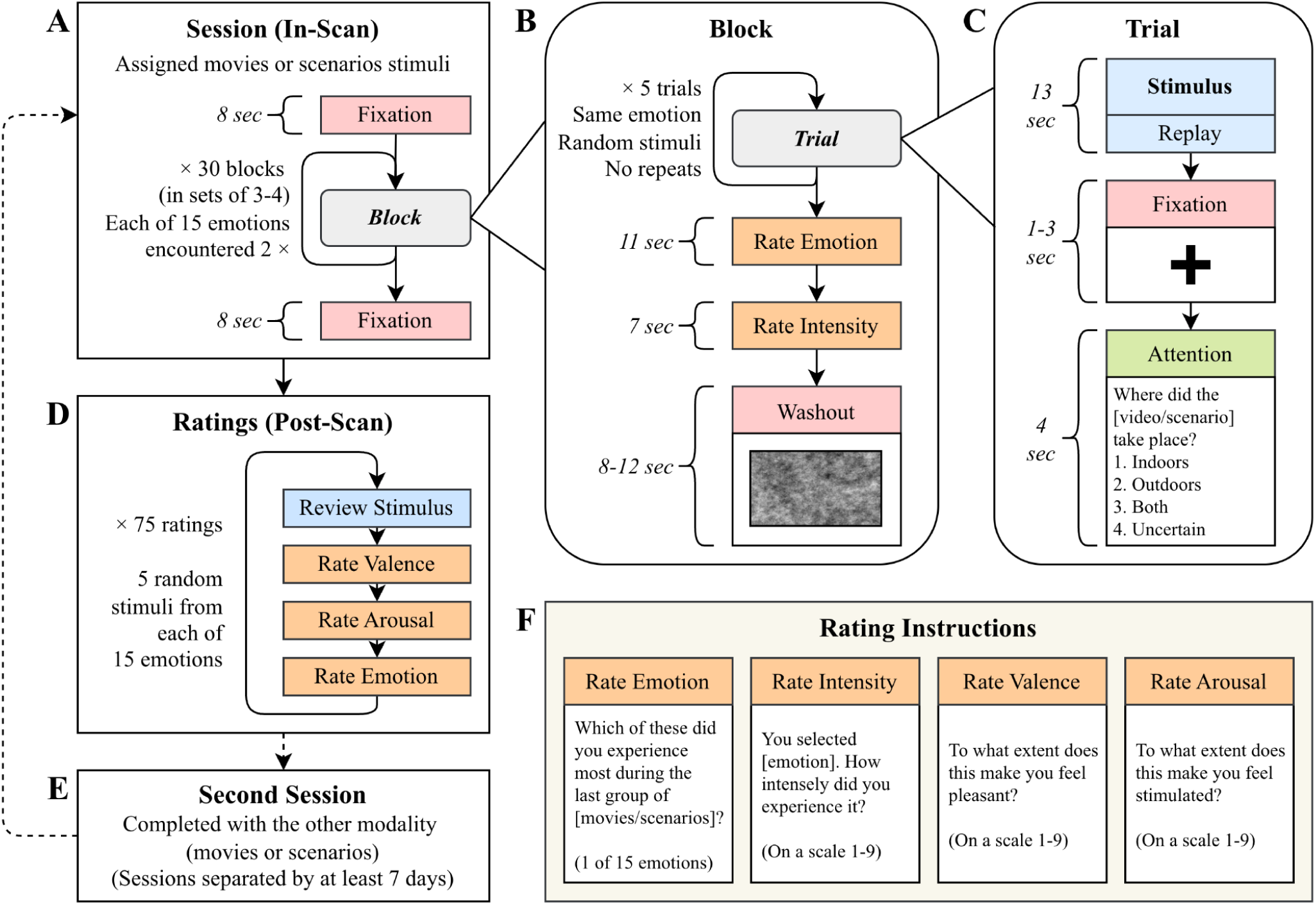
Task sequence flowchart. Each scanning session (A) is composed of thirty blocks (B), with each block containing five trials (C). (D) At the end of each scanning session, participants completed post-scan ratings for half of the 150 stimuli experienced during the session. (E) Participants completed two sessions, one with movie stimuli and one with scenario stimuli, in varying order. (F) Specific instructions used for collecting ratings.

#### MR imaging protocol

The MR imaging protocol at each session consisted of: a localizer acquisition, a high-resolution T1-weighted acquisition, a reverse-phase-encode-direction (RPED) echo planar imaging (EPI) acquisition, nine EPI acquisitions (8 task runs and a resting-state run), and another RPED EPI acquisition. The T1-weighted acquisition had the following parameters: acquisition matrix = 256 × 256, repetition time = 2250 ms, echo time = 3.12 ms, field of view = 256 mm, in-plane voxel size = 1.0 × 1.0 mm, slice thickness = 1.0 mm, spacing between slices = 0 mm, 192 axial AC-PC aligned slices. EPI acquisitions had the following parameters: acquisition matrix = 128 x 128, repetition time = 2000 ms, echo time = 30 ms, field of view = 256 × 256 mm, in-plane voxel size = 2.0 × 2.0 mm, slice thickness = 2.0 mm, spacing between slices = 0 mm, phase-encode direction = anterior-to-posterior, 69 axial slices angled 30 degrees positive of AC-PC, multiband acceleration factor = 3, in-plane acceleration factor = 2. Runs 1, 2, 3, 5, 6, and 7 of the functional task had 262 time-points (8 minutes, 44 seconds). Runs 4 and 8 had 200 time-points (6 minutes, 40 seconds). The resting state acquisition had 240 time-points (8 minutes). The RPED EPI acquisitions used the same parameters as the functional task EPI acquisitions except a phase-encode direction of posterior-to-anterior, and 2 time points.

### fMRI processing

#### fMRI preprocessing

Preprocessing and modeling of fMRI data were conducted with the following software suites: FreeSurfer v7.2.0 (42), fMRIPrep v23.0.2 (43), MRIQC v23.1.0 (44), dcm2niix v1.0.20211006 (45), Convert 3D v1.1.0 (c3d; 46), FMRIB Software Library v6.0.6.4 (FSL; 47), and Analysis of Functional NeuroImages v24.2.01 (AFNI; 48).

Anatomic T1-weighted and functional T2*-weighted DICOMs were converted to NIfTI format. FreeSurfer was used to generate anatomical parcellations from structural images. FMRIPrep subsequently integrated these outputs, the T2*-weighted functional scans, and the T2*-weighted RPED EPIs (for field inhomogeneity distortion correction) to produce preprocessed fMRI data warped into MNI152 template space; see Supporting Information, “FMRIPrep boilerplate” for complete details on the preprocessing pipeline. Next, additional preprocessing was conducted on the output of fMRIPrep to prepare the fMRI data for first-level modeling. These additional preprocessing steps included: highpass temporal filtering of the functional data (cutoff = 100 s; no lowpass filter), smoothing the data with a 6-mm FWHM 3D gaussian kernel, and scaling the output, such that all fMRI data had a common voxel intensity range.

#### fMRI modeling

First-level modeling of the functional data was performed using FSL’s FEAT (49), which fit a regression model (general linear model; GLM) to each fMRI run. The GLM included nuisance regressors for average CSF signal, average WM signal, framewise displacement, and motion correction translation and rotation parameters and their derivatives. Motion-censoring regressors (values of 0 at all time-points except for one, which had a value of 1) were also included for time-points with framewise displacement values greater than 0.5 mm. Task-related regressors were created by convolving boxcar functions of each task component’s on/off timing with a double-gamma hemodynamic response function. These task components included induction stimulus presentation, mental replay instructions, the location judgement prompt, emotion endorsement, emotion degree rating, and washout picture presentation. Importantly, stimulus regressors were created on a block-wise basis in order to obtain activation contrast estimates for every block individually. For each block, an Induction Stimulus Block > Washout contrast was generated for use as input to the multivariate regression analysis. After first-level modeling, a gray-matter mask including cerebellar, subcortical, and brainstem gray matter was applied to the whole-brain emotion vs. washout contrast maps. After masking, the number of voxels in each contrast map was 158,110.

### Multivariate pattern analysis

#### Model selection

Ridge, least absolute shrinkage and selection operator with principal component regression (LASSO-PCR), Elastic Net with principal component regression (ElasticNet-PCR), partial least squares regression (PLS), and linear support vector regression (SVR-Linear) were used for multivoxel pattern analysis. Only linear models were implemented, considering they are straightforward to interpret, and have been found to perform as well, if not better than, non-linear models at decoding fMRI data (15,21). Analysis was performed across subjects, rather than within, to capture group-level neural patterns associated with arousal and valence and poise conclusions for generalization to new individuals as opposed to individual differences. When using voxel-wise contrast values to decode arousal and valence, the number of features (158,110 voxels) greatly exceeds the number of observations (nearly 3,200 blocks). Additionally, high multicollinearity can be expected among features considering the inherent spatial autocorrelation of fMRI data and the shared variance between co-activated regions of the brain. MVPA models were selected to address these characteristics.

Penalized regression models, including LASSO, Ridge and ElasticNet, all effectively handle large numbers of features relative to observations (50). However, voxel selection by models with an L1 penalty, like LASSO and by extension ElasticNet, can be unstable; neighboring voxels have high correlation and these models will arbitrarily select a few features from correlated groups (51). To maintain voxel weight map stability, principal component analysis (PCA) was applied prior to LASSO and ElasticNet. Known as principal component regression (PCR), this approach allowed models to act on orthogonal components with reduced covariation (52) and has previously found success in emotion decoding of fMRI data (13).

We included linear support vector regression (SVR) based on the previous success and common use of support vector machines in fMRI decoding (14,15). Partial least squares regression (PLS) has also been used with success in the decoding of emotion-related data using fMRI (53). This method fits a linear regression model within a lower-dimensional subspace constructed to maximize the covariance between features and targets and is well-suited for cases with more features than observations and multicollinearity among features (54).

The multivariate analysis is depicted in Fig 8. While not the primary focus of this study, a parametric modulation analysis was also included to complement the multivariate decoding. The parametric modulation analysis is univariate and identifies voxels whose BOLD signal scales with either trial-wise arousal (from low to high) or valence (from negative to positive) levels. A full description of the parametric modulation analysis and results can be found in Supporting Information “Parametric modulation analysis.”

**Fig 8.**
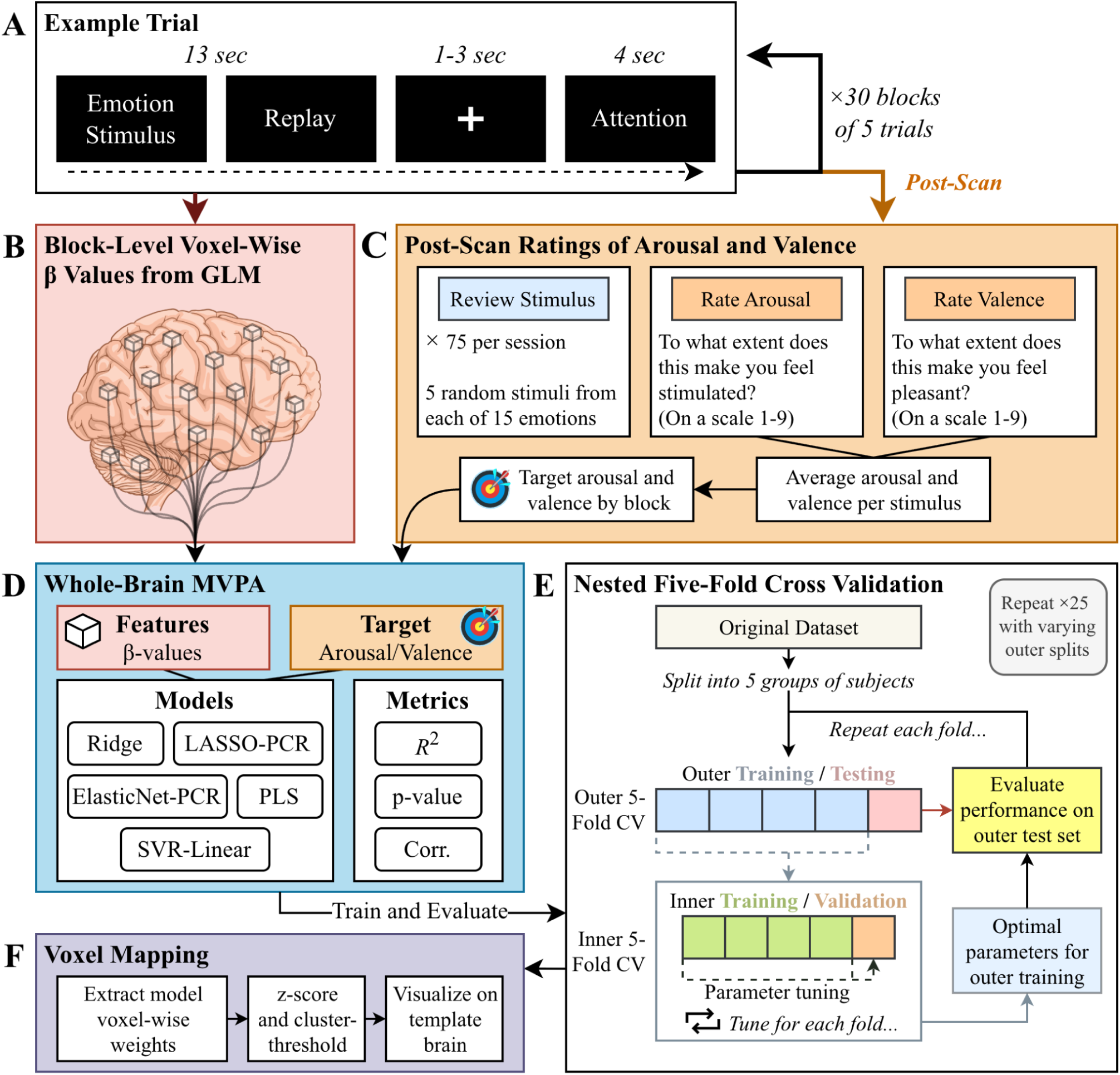
Analysis flowchart. (A) Participants experienced 30 blocks of five trials of affective movies and scenarios. See Fig 7 for more details regarding the task. (B) Mass univariate GLMs identified whole-brain voxel-wise β values at the block-level. The brain illustration is adapted from NIAID Visual & Medical Arts (55). (C) Participants rated stimuli for arousal and valence following the scan, and these values were used as targets in the multivariate pattern analysis (MVPA). (D) Whole-brain MVPA was conducted using a number of models and evaluated with a variety of metrics, such as *R*^2^, *p*-value, and Pearson’s correlation (Corr.). (E) MVPA models were trained using a nested five-fold subject-independent cross-validation (CV). The training of each model was repeated 25 times with varying training and testing splits to increase power and stabilize weight maps. (F) Important voxel-wise coefficients were extracted from the MVPA models and mapped onto the MNI152 template brain for examination of relevant regions. Abbreviations: ElasticNet-PCR = elastic net with principal component regression; LASSO-PCR = least absolute shrinkage and selection operator with principal component regression; PLS = partial least squares regression; SVR-Linear = linear support vector regression.

#### Targets

The targets in the MVPA were block-level arousal and valence values. To calculate these measures, the mean arousal and valence was first calculated for each stimulus using post-scan ratings from all participants who completed sessions, regardless of whether their fMRI data passed quality control. Next, the arousal and valence values for each block were calculated as the mean of the block’s five stimulus’ arousal and valence values. As opposed to taking the mean of all ratings across stimuli directly, which would weight stimuli by the number of raters, this approach was chosen to reflect the experimental structure, in which each stimulus was presented for equal duration and thus approximately contributed equally to the participant’s experienced affective state. The MVPA was conducted at the block level, with these block-level arousal and valence values acting as targets. These targets represent the expected arousal/valence state of a participant experiencing that block.

#### Model training and testing

All multivariate regression analyses and associated metric calculations were completed with SciKit-Learn v1.6.1 (56). A five-fold subject-independent cross-validation was used, where data from 80% of the participants were used to train the regression models, and the remaining 20% of participants were used to test performance until all participants had been used for testing (see Fig 8E). The subject-independent nature of the cross-validation ensures independence between the train and test sets considering blocks from one participant may correlate. Within the training data, a second five-fold subject-independent inner loop was used to set the model parameters. This nested cross-validation scheme limits optimistic biases (51). More details about the parameter tuning can be found in Supporting Information, “Model parameter tuning.” Since larger ranges can disproportionately influence certain models’ behavior, the features were also z-scored before analysis to equalize potential influence. To prevent data leakage, the training and test sets were scaled independently using a SciKit-Learn Pipeline (56). Considering that the performance of a model can be overestimated from a single cross-validation split due to the randomness of the partitions, each model was repeated 25 times with randomized inner and outer training and testing splits to generate a more representative distribution of performance, increase power, and stabilize weight maps (51,57,58). Performance was measured by calculating the *R*^2^, Pearson’s correlation, and root-mean-squared error between the model’s predictions and the targets.

Significance was estimated for each model via permutation testing (59). Each model was repeated 1000 times with targets permuted, breaking any relationship to the fMRI features so chance performance can be estimated. Given that each participant is shown 30 blocks that sample the arousal-valence space in a randomized sequence, to satisfy the assumption of exchangeability, we permuted target values within each participant (58). Permutation test repetitions were run the same as with the true labels, using a five-fold subject-independent inner and outer cross-validation scheme with identical parameter tuning (57,58). *P*-values were estimated empirically by calculating 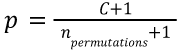, where *C* is the number of permuted models that out-perform the true score, and *n_permutations_* is the total number of permutations (56,59). The number of permutations was set to 1000, producing a minimum *p*-value of 0.000999. Approximate computational costs for each model are described in Supporting Information, “MVPA computational efficiency” and may assist MVPA model selection in future research, in addition to aiding in reproducibility.

#### Voxel mapping

The coefficients of linear models represent the impact a feature has on the estimation of the target. To visualize which voxels and regions are most important in predicting arousal and valence, the model coefficients were averaged across all folds of all repetitions and z-scored. Coefficients of PCR models were first mapped from component space back to voxel space by multiplying the loadings of the PCA components with the coefficients of the model (52). Considering that coefficients could be positive or negative, a two-tailed threshold corresponding to p < 0.05 (|z| > 1.96) was applied to identify significant voxels. A threshold of p < 0.05 is consistent with past affect-related whole-brain decoding studies (17,19,53). Additionally, to reduce false positives, only clusters having 30 or greater voxels, defined using first nearest-neighbor connectivity (voxels sharing a face), are included in visualizations. Clustering was conducted using the cluster command in FSL v.6.0.6.4 (47). Anatomical labels were assigned using the Harvard-Oxford Cortical and Subcortical Structural Atlases, and the Cerebellar Atlas in MNI152 Space after Normalization with FNIRT, all of which are included with FSL v6.0.6.4 (47). The Harvard Ascending Arousal Network Atlas v2.0 (60) was additionally used in assigning labels to brainstem regions. Maps were visualized using MRIcroGL v1.2.20220720 (27,61). With these settings, the resultant maps are expected to accurately indicate which regions’ activation or deactivation, taken together, reliably predict arousal or valence.

## Data and code availability

Code for all steps of the analysis is publicly available at https://github.com/labarlab-emorep-AVDecoding. Particularly, code used to decode arousal and valence is found at https://github.com/labarlab-emorep-AVDecoding/av_regression. Additionally, this repository contains complete cluster tables and visualizations for all five implemented MVPA models for arousal and valence across both induction modalities. Multivariate pattern analysis was completed with SciKit-Learn v1.6.1 (56). The raw data is publicly available at https://nda.nih.gov/edit_collection.html?id=3639. The REFORMS checklist (62) was used when preparing this manuscript.

## Acknowledgements

The authors thank Mary Baumann, Shannon Chen, Sebastian Sanchez, Alisa Schutz, and Cody Triplett for their assistance with participant recruitment and data collection.

