## Supporting Information for "Decoding of arousal and valence from fMRI data obtained during emotion inductions"

### Full task description

#### MRI day schedule

Participants completed the BDI-II and MR safety screening forms the morning of each MRI session before arrival. Sessions were canceled if participants scored 20 or higher on the BDI-II, as they no longer met study inclusion/exclusion criteria. Upon arrival, participants completed the Positive and Negative Affect Schedule (PANAS) and the Spielberger State-Trait Anxiety Inventory (STAI) questionnaires. They were then oriented to the logistics of the MRI scan and trained on the functional imaging task. A practice version of the task was also administered by study personnel to ensure that participants understood and could perform the task. Skin conductance electrodes were then put on the participant's hand and the MR technologist helped them into the MRI scanner bore.

#### fMRI task hardware

A Siemens Magnetom 3.0T scanner was used for MR imaging. A MRI-compatible 4K monitor (InroomViewingDevice, NordicNeurolabs, Bergen, Norway) at the rear of the scanner presented the functional task to participants via a mirror attached to the head coil. Participants responded to the functional task with a 4-button response box in the right hand (Current Designs, Inc., Philadelphia, PA).

#### Physiology recording hardware

Although not leveraged in the present study, three physiology signals (pulse, respiration, and galvanic skin response (GSR)) were collected with a BIOPAC MP160 (BIOPAC Systems, Inc., Goleta, CA) while participants performed the functional tasks in the MR scanner. These were recorded from, respectively, a MR-compatible finger pulse oximeter, a respiration belt, and GSR hand electrodes. Before entering the scanner room, participants had two GSR electrodes placed on the hypothenar eminence of the left hand, perpendicular to the eminence. The finger pulse oximeter was positioned around the index finger of the left hand, and the respiration belt was placed around the torso. Recording of physiological signals was time-matched to the functional emotion induction task via a "code" channel in AcqKnowledge, which read value changes passed from the task code to the BIOPAC.

#### Emotion induction task

Participants performed an emotion induction task while undergoing fMRI scanning. This task consisted of six parts: passively viewing emotional videos or reading emotional text scenarios; mentally replaying the most recent stimulus; answering a prompt about where the video or scenario took place (indoor/outdoor); reporting which emotion (from a list of 15) was experienced the most during the preceding block of stimuli; reporting to what degree the reported emotion was experienced; passively viewing grayscale "washout" pictures.

Participants completed the task over two sessions, separated by at least seven days. The two sessions were identical except for the type of induction stimulus; movie stimuli were used one day and scenario stimuli were used the other day. The order of stimulus type was balanced across participants. Each session had eight task runs. These runs had a total of 30 task blocks, with each run having 3 or 4 blocks. The beginning and ending of each run consisted of an 8-second presentation of a small black cross in the center of the screen. Each task block consisted of 5 emotion induction trials followed by the emotion endorsement and degree experienced questions and, finally, passive viewing of a washout picture.

Videos were presented without sound and were followed by the instruction “Replay the video in your mind”. The duration of the “replay” instruction was tailored for each video such that the total length of the video plus the “replay” time equaled 13 seconds. Text scenarios were presented for a duration equal to the average video length of the associated emotion induction category. Similarly, each scenario presentation was followed by the instruction “Replay the scenario in your mind” and total stimulus and replay duration was set to 13 seconds.

A two-second (average, jittered 1-3 seconds) fixation cross was displayed after each “replay” instruction. A question about where the events of the stimulus took place was then presented: “Where did the [video/scenario] take place?”; possible responses were “1: Indoors”, “2: Outdoors”, “3: Both”, and “4: Uncertain”. Participants had 4 seconds to select one of these options using a button-box held in the right hand. These questions were included as attention checks.

Following the final (5th) trial of each block, participants were asked to endorse the emotion experienced the most during that block. A list of 15 emotions (amusement, anger, anxiety, awe, calmness, craving, disgust, excitement, fear, horror, joy, neutral, romance, sadness, surprise) was shown down the center of the screen along with the prompt “Which of these did you experience the most during the last group of [movies/scenarios]?” The list order was randomized for every presentation. Participants used two buttons on the button box to move a selection cursor up or down, starting from the center of the list, and had 11 seconds to move the cursor to the emotion they wished to select. After the emotion endorsement, participants reported to what degree they experienced the endorsed emotion via a similar screen. Integers 1 through 10 were displayed as a list in the center of the screen and participants again moved a selection cursor up or down to select a response, starting from the bottom of the list. The message “You selected [emotion]. How intensely did you experience it?” was also displayed on this screen and participants had 7 seconds to move the cursor to their desired integer.

Finally, after the emotion intensity rating, one of three grayscale “washout” pictures was displayed for 10 seconds (average, jittered 8-12 seconds). These pictures were created from three neutral pictures (of flowers and trains) which were converted to grayscale and then phase-scrambled. The final pictures retained some low-level characteristics of the originals, but contained no discernable objects.

##### Post-scan arousal and valence ratings
Following the scanning session, participants rated the arousal and valence of a random 75 of the total 150 stimuli shown during the scanning session. To rate valence, participants were asked “To what extent does this make you feel pleasant?” with integers on a scale from 1 to 9, where 1 is “very unpleasant” and 9 is “very pleasant”. To rate arousal, participants were asked “To what extent does this make you feel stimulated?” with integers on a scale from 1 to 9, where 1 is “very subdued” and 9 is “more stimulated.”

#### Parametric modulation analysis
A parametric modulation analysis was conducted on the functional imaging data to identify regions of the brain in which blood-oxygenation-level-dependent (BOLD) signal change was related to either changes in arousal or changes in valence. A general linear model (GLM) was constructed for each run containing the following

task-related regressors: stimulus presentation, modulated by arousal; stimulus presentation, modulated by valence; mental replay, modulated by arousal; mental replay, modulated by valence; all stimulus presentations; all mental replay sections; location judgement prompt; emotion endorsement; emotion degree rating; and washout picture presentation. As with the GLM for the primary analysis, these regressors consisted of boxcar functions convolved with a double-gamma hemodynamic response function. For the modulated regressors, the relative height of each "on" portion of the boxcar function was scaled by either the arousal rating or the valence rating of the associated stimulus being presented. The arousal and valence ratings for each stimulus were constructed as the group averages from the post-scan ratings. These values were then mean-centered to create the final modulation weights.

The parametric modulation GLM contained the same nuisance regressors as that of the primary analysis (see "fMRI Modeling" section for details): average CSF, average WM, framewise displacement, motion correction and derivatives, and motion-censoring regressors.

Run-level analyses were entered into session-level analyses, creating whole-brain GLM beta maps for each session. These were then combined within stimulus type (movies, scenarios) across participants for group-level analyses. For multiple-comparison correction, final group analyses used a voxel-wise z-score threshold of 3.1 and a cluster-wise alpha $< 0.05$.

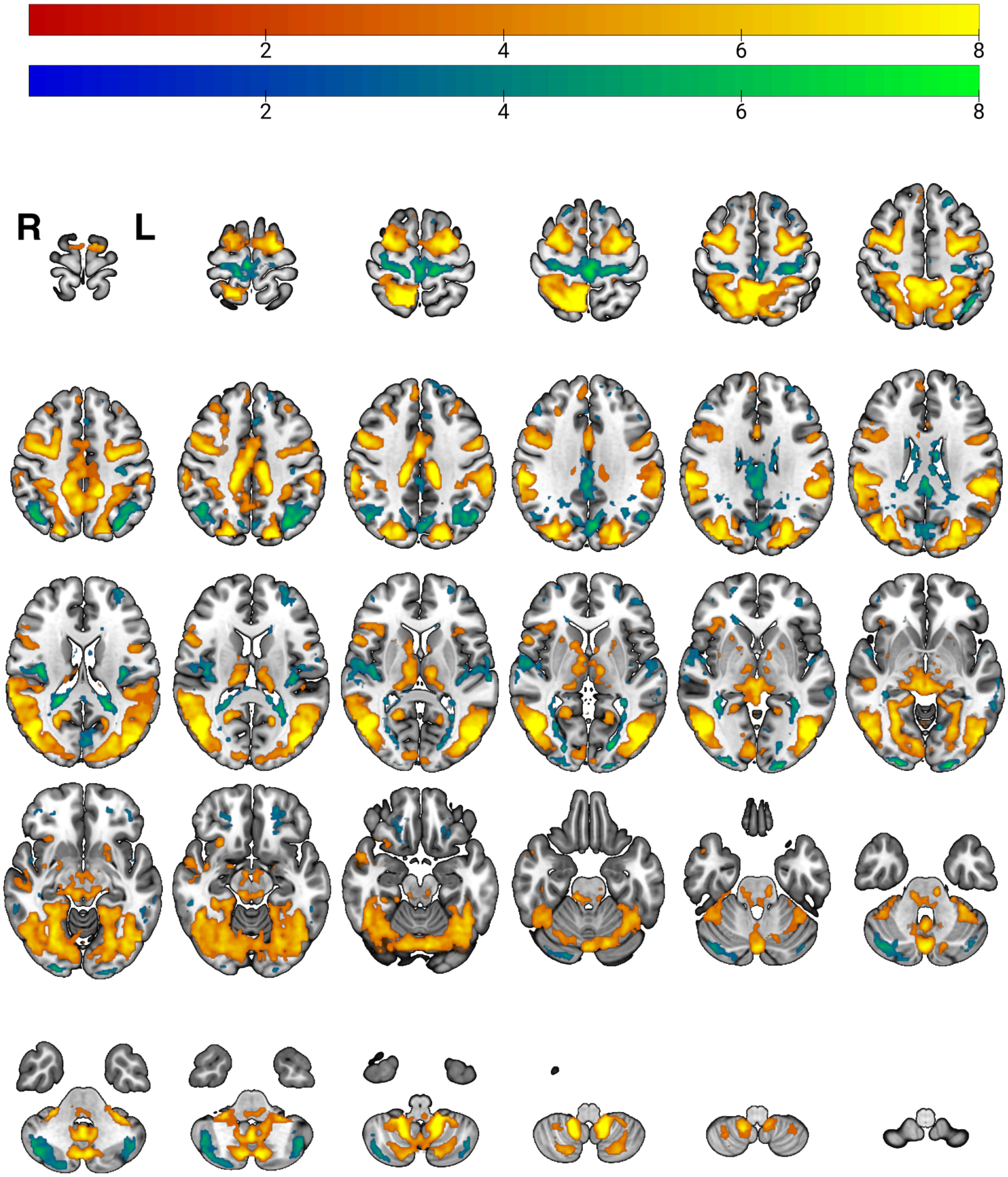


**Fig S1. Evenly spaced axial slices showing significant clusters for the parametric modulation of arousal during movie viewing.**

Colored overlays indicate regions where activity varied with arousal ratings. Negative coefficients are displayed in blue-green, positive in red-yellow.

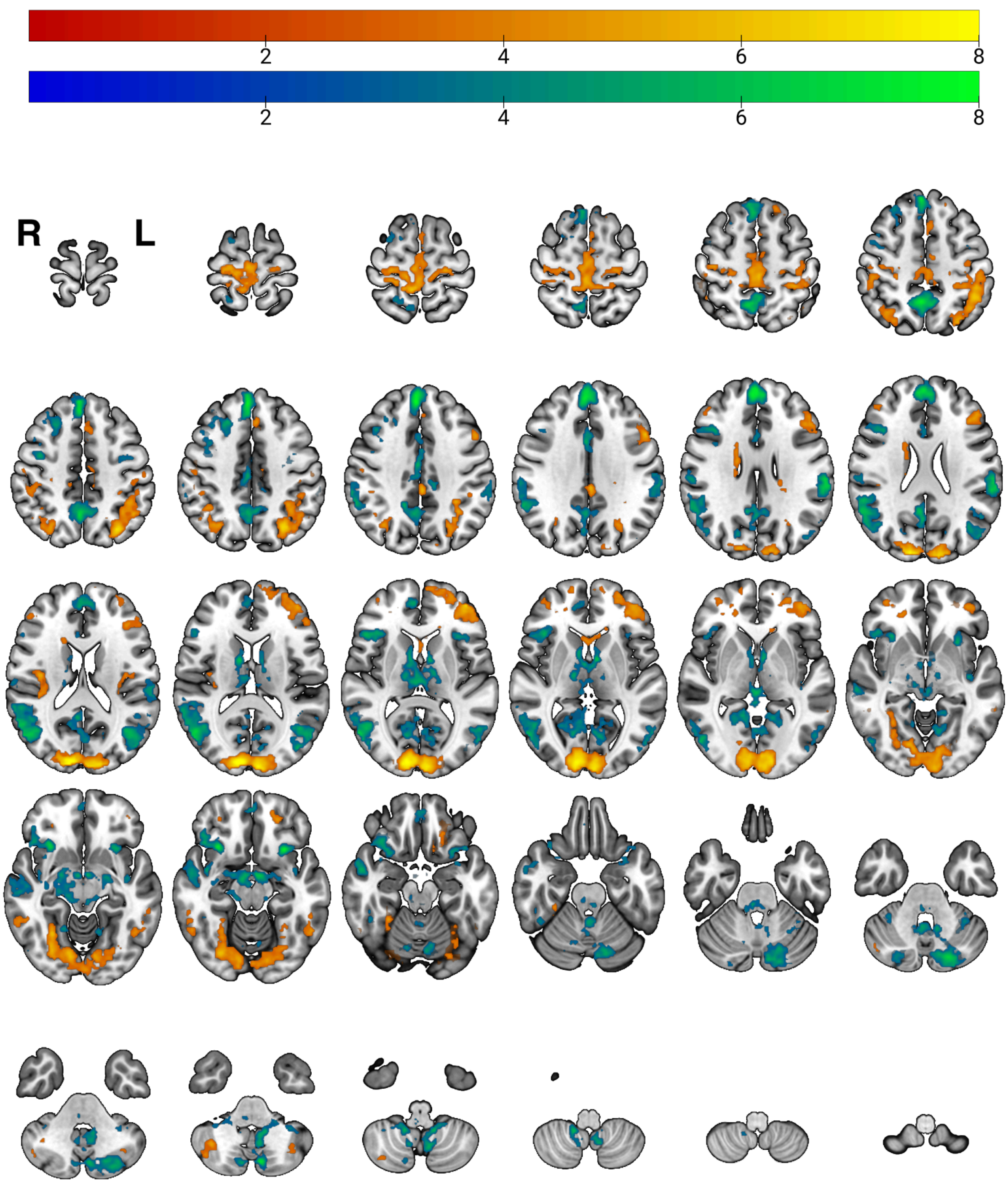


**Fig S2. Evenly spaced axial slices showing significant clusters for the parametric modulation of valence during movie viewing.**

Colored overlays indicate regions where activity varied with valence ratings. Negative coefficients are displayed in blue-green, positive in red-yellow.

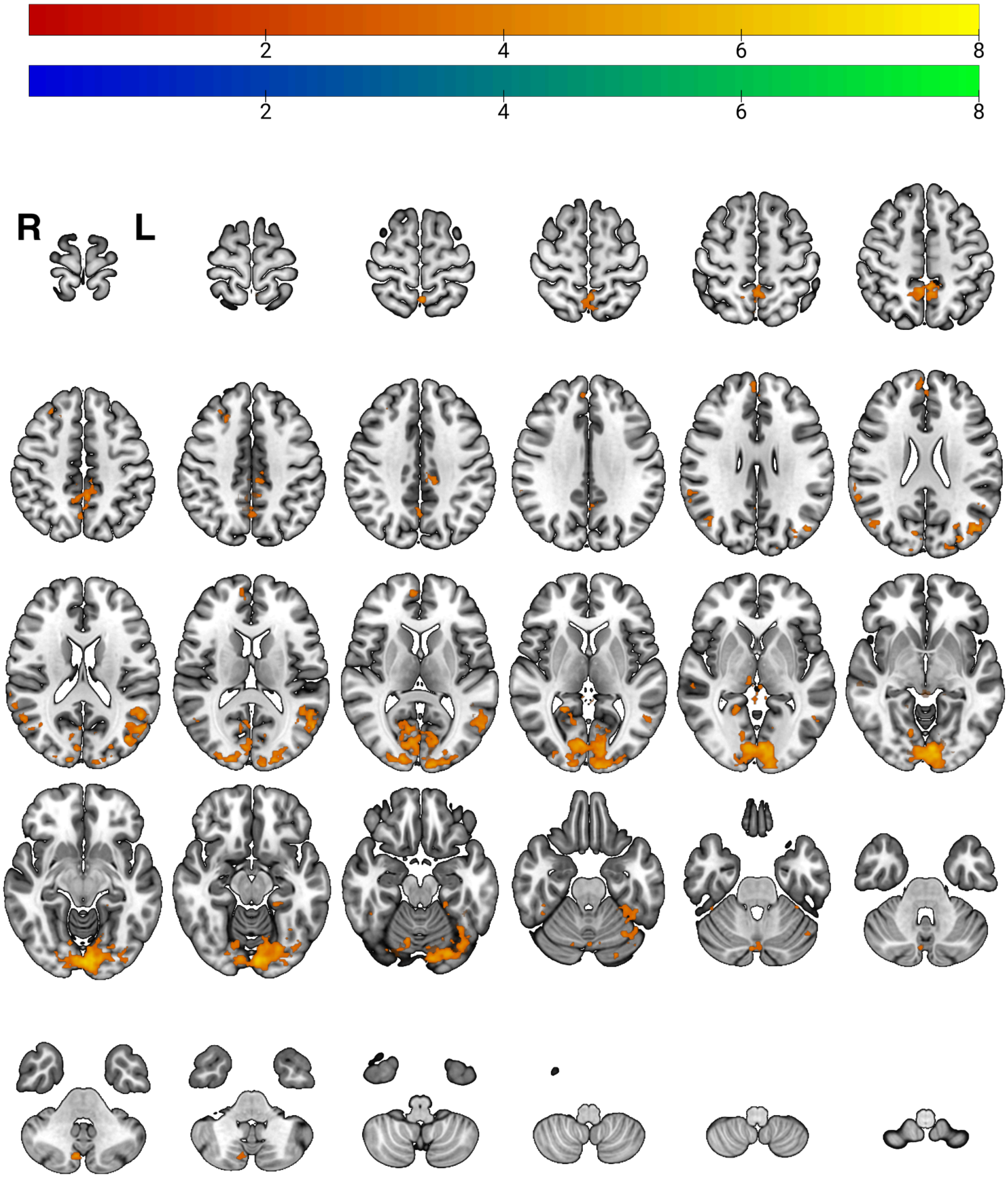


**Fig S3. Evenly spaced axial slices showing significant clusters for the parametric modulation of arousal during scenario viewing.**

Colored overlays indicate regions where activity varied with arousal ratings. Negative coefficients are displayed in blue-green, positive in red-yellow.

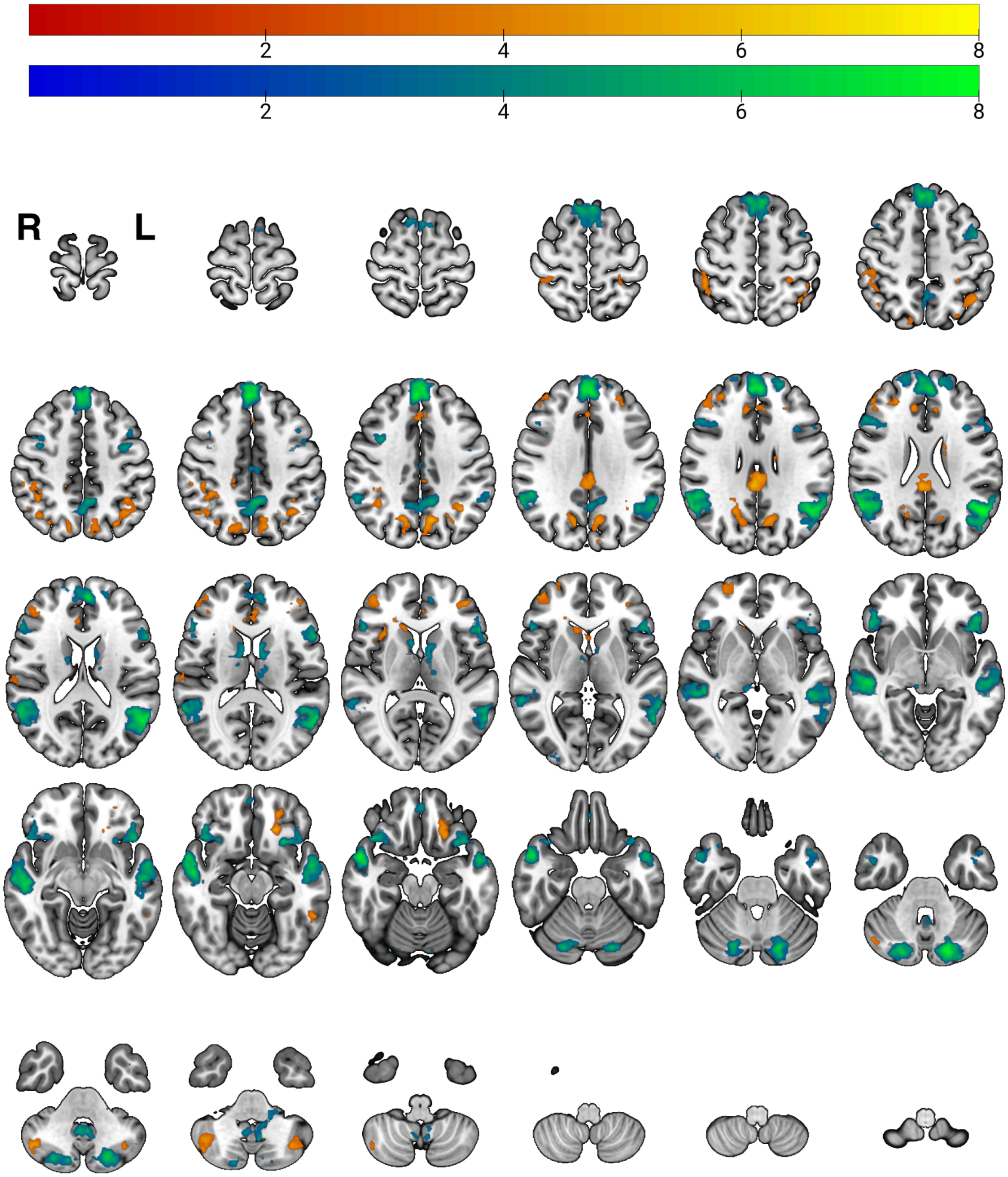


**Fig S4. Evenly spaced axial slices showing significant clusters for the parametric modulation of valence during scenario viewing.**

Colored overlays indicate regions where activity varied with valence ratings. Negative coefficients are displayed in blue-green, positive in red-yellow.

#### Movie and scenario stimuli

Ten clips were selected for each emotion category using published normative ratings (1) supplemented with our own ratings for clips not present in that dataset. Additional clips were added to better populate categories that were underrepresented in the original norms (e.g., anger, calmness, excitement, neutral). Following procedures similar to Cowen & Keltner (1) and Faul et al. (2), we obtained normative judgments from participants who chose the emotion label(s) (from a set of 34 options; boredom was relabeled as neutral) that best matched their experience while viewing each clip. Each clip was rated by at least nine participants in Cowen & Keltner (1) and by at least 11 participants in our supplementary norming. Clips were retained only if the target category achieved a concordance (agreement) rate > 0.5 and exceeded the concordance for any other category. Clips ranged from 3–12 s (M = 6.8 s, SD = 2.3 s). Although a one-way ANOVA across the 15 emotion categories indicated a small overall difference in mean clip duration ($F(14, 135) = 1.99$, $p = 0.023$), no pairwise contrasts survived Tukey correction (all $p > 0.05$); only marginal shorter duration for romance versus several categories was observed. The mean normative concordance for movies was 0.73 (SD = 0.13), with disgust showing the highest mean agreement (0.97) and anxiety and excitement among the lowest (0.62). Concordance rates differed significantly across categories ($F(14, 135) = 16.89$, $p < .001$); post hoc tests indicated that disgust exceeded most other categories (except amusement), while several categories (e.g., amusement, sadness, craving, fear, calmness) also showed relatively high agreement and others (e.g., anxiety, excitement, awe) showed comparatively lower agreement.

Scenario stimuli (ten per category) were drawn from the norming reported in Faul et al. (2), which includes both newly developed vignettes and items adapted from Fields & Kuperberg (3). As in the movie norming, MTurk participants (≥11 raters per scenario) selected the emotion category that best described how they would feel when imagining each scenario; items were accepted only when the intended category's concordance exceeded 0.5 and surpassed other categories. All scenarios were written in the second person and consisted of two sentences (mean words = 19.31, SD = 4.52). One-way ANOVAs showed no reliable differences across the 15 categories in word count ($F(14, 135) = 1.53$, $p = 0.108$), text concreteness ($F(14, 135) = 1.25$, $p = 0.245$), or imageability ($F(14, 135) = 0.79$, $p = 0.682$); concreteness and imageability were quantified using TAALES 2.0 metrics (4,5) and the MRC database (6). The average concordance for scenarios was 0.75 (SD = 0.09); fear had the highest mean agreement (0.90) and neutral the lowest (0.66). Concordance varied by category ($F(14, 135) = 8.45$, $p < .001$). Tukey HSD follow-up tests indicated that fear elicited higher concordance than all other categories except disgust (all $p < .05$). Disgust also showed higher concordance than most other categories, whereas neutral and awe exhibited comparatively lower concordance than several categories.

#### Model parameter tuning

All parameters were optimized for each outer fold using an inner subject-independent 5-fold cross-validation. LASSO-PCR and ElasticNet-PCR were optimized for mean squared error from a range of 100 values generated automatically, the default in SciKit-Learn (7). ElasticNet-PCR was tuned for 100 alphas at each of seven different ratios of L1 to L2 regularization. Ridge was optimized for $R^2$, the default regression cross-validation in SciKit-Learn (7), across a logarithmic scale of 50 alpha values. PCA, which is used prior to fitting in ElasticNet-PCR and LASSO-PCR, reduced the data to $n$ components, where $n$ equals the minimum of the number of observations and the number of features minus 1, as is the default in SciKit-Learn. PLS was optimized for $R^2$ across a range of $n$ components from 1 to 20. Linear-SVR was optimized for $R^2$ across a logarithmic grid of C, 6 values from 1e-6 to 0.1, and epsilon, 8 values from 1e-2 to 1. Parameter ranges were selected to reach adequate performance while balancing computational efficiency.

MVPA parameter grids are often manually refined based on observation and exploratory tuning. To improve transparency and leverage a larger empirical basis, we instead implemented an automated procedure to tighten the parameter grids for PLS, Ridge, and Linear-SVR by observing the top parameters. After an initial inspection phase, we examined which parameter values were selected across folds in the 25 true-target repetitions and in the first number of permutations, and removed values that were never chosen. 100 permutations were used for PLS and Ridge, and 25 with SVR-Linear considering its computational cost. To avoid over-pruning, we extended the observed selection range by a small margin, more specifically retaining a number of parameters nearby to any that were selected, and, for PLS and SVR-Linear which reported metrics for each parameter tested in each search, retained any parameter value within 0.01 $R^2$ of the best-performing values. This procedure substantially reduced compute time for the full permutation set. The implementation of this procedure is provided in the project repository (see Reproducibility). As aforementioned, parameters in LASSO-PCR and ElasticNet-PCR were optimized for mean squared error automatically, eliminating the need for parameter grid reduction.

#### MVPA computational efficiency

Computational cost for each decoding model was recorded during execution on a shared high-performance cluster. For each recorded model repetition, we logged wall-clock time and requested resources. We computed normalized CPU cost as wall clock-hours per repetition multiplied by the number of CPUs requested and summarize as median ± IQR across available repetitions. Similarly, GB-hours were calculated as wall clock-hours per repetition multiplied by the RAM requested in gigabytes. Because the cluster was a shared resource and jobs were not isolated to dedicated nodes, runtimes were influenced by variable I/O and load; all timings should therefore be interpreted as approximate cost estimates for planning and reproducibility. Additionally, computational cost depends critically on the size and complexity of the model's parameter search, with more exhaustive searches substantially increasing runtime and resource usage. See Supporting Information, "Model parameter tuning" for more details regarding model parameter tuning. Resource allocations aimed to maximize efficiency and resource usage while minimizing the resources requested. Model parameter searches and fitting were parallelized across all CPUs if possible. Although performance can be relatively similar between models, computational cost varies significantly. Table S1 details core-hours and GB-hours by model, and Fig S5 displays core-hours and GB-hours as a boxplot. Based on the median core-hours and GB-hours reported in Table S1, completing 1,000 permutations across all model-target-task combinations required approximately 344,810 core-hours and 3,136,720 GB-hours in total. Memory was the primary limiting factor due to the large input size (~3,100 blocks and 158,110 features).

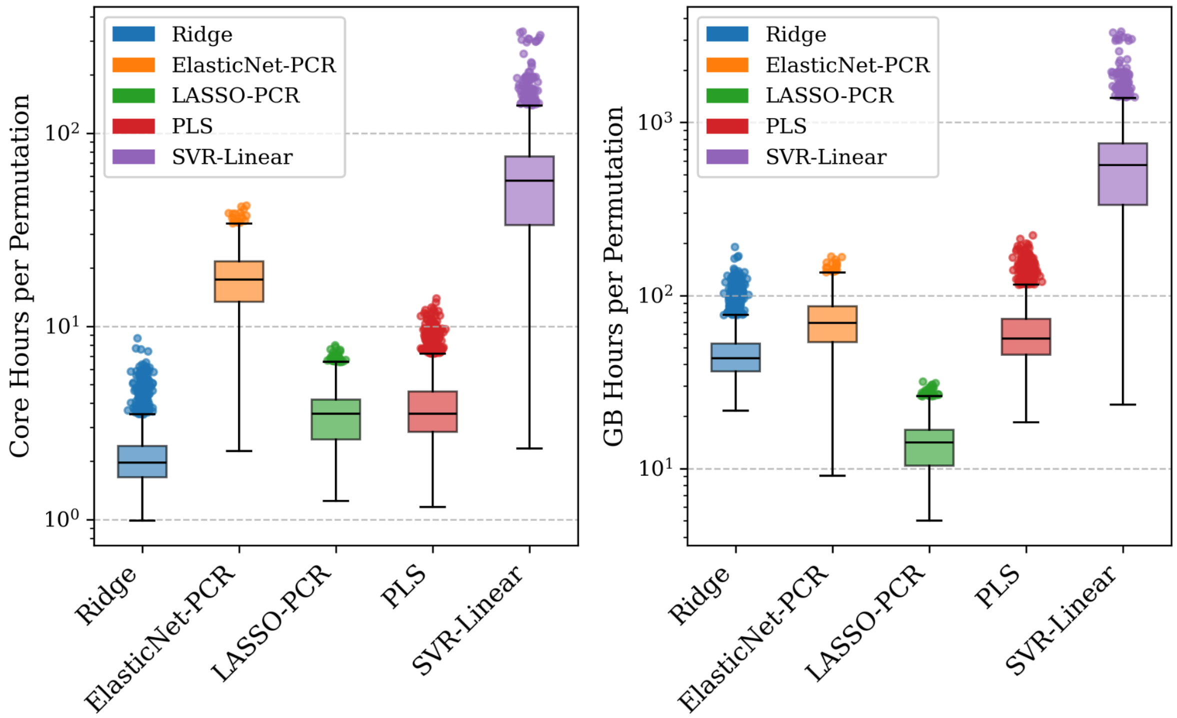


**Fig S5. Computational cost per model.**
Boxes show median ± IQR of core-hours per model; jittered points are individual repetitions. Core-hours were computed as wall-clock hours multiplied by the number of CPU cores requested (i.e. n_jobs). GB-hours were calculated as wall-clock hours multiplied by the requested memory in gigabytes. Runtimes were measured on a shared cluster and include system variability.

| Model | Target | Task | Random States Recorded | Requested Cores | Requested Memory (GB) | Median Core Hours | Core Hours IQR | Median GB Hours | GB Hours IQR |
|---|---|---|---|---|---|---|---|---|---|
| Ridge | Arousal | Movies | 868 | 1 | 22 | 1.66 | (1.55, 2.11) | 36.42 | (34.17, 46.41) |
| | | Scenarios | 800 | 1 | 22 | 1.72 | (1.42, 2.09) | 37.95 | (31.18, 46.05) |
| | Valence | Movies | 894 | 1 | 22 | 2.07 | (1.92, 2.53) | 45.53 | (42.29, 55.77) |
| | | Scenarios | 896 | 1 | 22 | 2.27 | (1.91, 2.91) | 49.96 | (42.09, 64.12) |
| ElasticNet-PCR | Arousal | Movies | 894 | 8 | 32 | 17.99 | (13.96, 22.07) | 71.97 | (55.84, 88.28) |
| | | Scenarios | 896 | 8 | 32 | 16.38 | (12.91, 21.29) | 65.51 | (51.63, 85.16) |
| | Valence | Movies | 918 | 8 | 32 | 16.81 | (12.96, 21.64) | 67.22 | (51.84, 86.54) |
| | | Scenarios | 899 | 8 | 32 | 17.99 | (14.53, 21.58) | 71.97 | (58.14, 86.33) |
| LASSO-PCR | Arousal | Movies | 891 | 8 | 32 | 2.76 | (2.17, 3.71) | 11.05 | (8.69, 14.85) |
| | | Scenarios | 900 | 8 | 32 | 3.51 | (2.61, 4.3) | 14.04 | (10.44, 17.19) |
| | Valence | Movies | 896 | 8 | 32 | 3.67 | (2.53, 4.27) | 14.66 | (10.12, 17.07) |
| | | Scenarios | 894 | 8 | 32 | 3.81 | (3.28, 4.37) | 15.22 | (13.11, 17.49) |
| PLS | Arousal | Movies | 895 | 6 | 96 | 2.52 | (2.06, 3.08) | 40.29 | (33.01, 49.2) |
| | | Scenarios | 800 | 6 | 96 | 3.5 | (3.26, 3.71) | 55.94 | (52.21, 59.33) |
| | Valence | Movies | 898 | 6 | 96 | 5.92 | (3.84, 6.92) | 94.73 | (61.43, 110.72) |
| | | Scenarios | 900 | 6 | 96 | 3.66 | (3.37, 4.46) | 58.53 | (53.91, 71.29) |
| SVR-Linear | Arousal | Movies | 860 | 16 | 160 | 95.63 | (63.21, 114.81) | 956.35 | (632.12, 1148.12) |
| | | Scenarios | 998 | 16 | 160 | 59.15 | (40.96, 70.7) | 591.47 | (409.57, 706.95) |
| | Valence | Movies | 943 | 16 | 160 | 61.74 | (41.91, 72.46) | 617.39 | (419.13, 724.63) |
| | | Scenarios | 810 | 16 | 160 | 22.05 | (14.8, 28.01) | 220.52 | (148.01, 280.1) |

**Table S1. Summary of computational costs for null permutations of decoding models.**
Each row shows a unique combination of model, target, and task. Reported values include the number of random states recorded, requested CPU cores and memory per job, and normalized runtime metrics: core hours being wall-clock hours multiplied with requested cores, and GB hours being wall-clock hours multiplied with requested memory in gigabytes. Medians and intraquartile range (IQR) summarize variability across available repetitions. Runtimes were measured on a shared HPC cluster. Peak memory was not recorded for all repetitions, so reported GB reflects requested memory.

### Model performance versus permutations

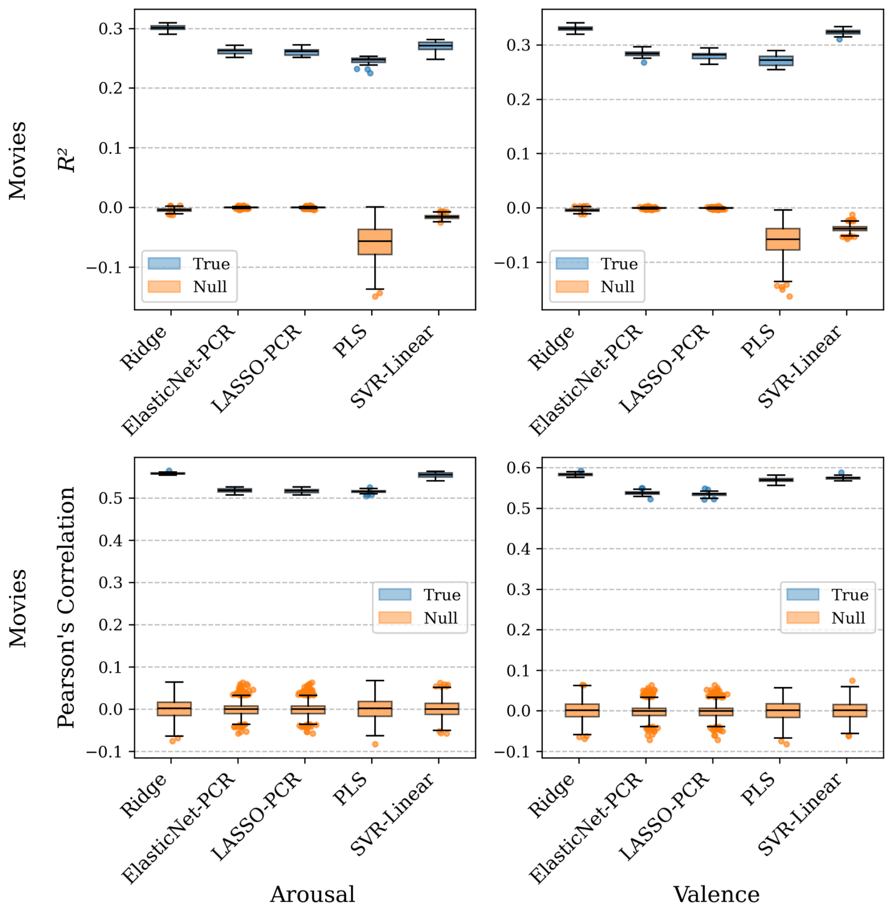


**Fig S6. Boxplots depicting all models' performance in decoding arousal (left) and valence (right) from fMRI data collected during movies viewing.**

The top row shows $R^2$ and the bottom row shows Pearson's correlation. The distribution of each models' 25 repetitions are shown in blue and their associated 1000 permutations, which were used as a baseline for calculating *p*-value, in orange. The permutation tests indicate that all are significant with a *p*-value less than 0.001.

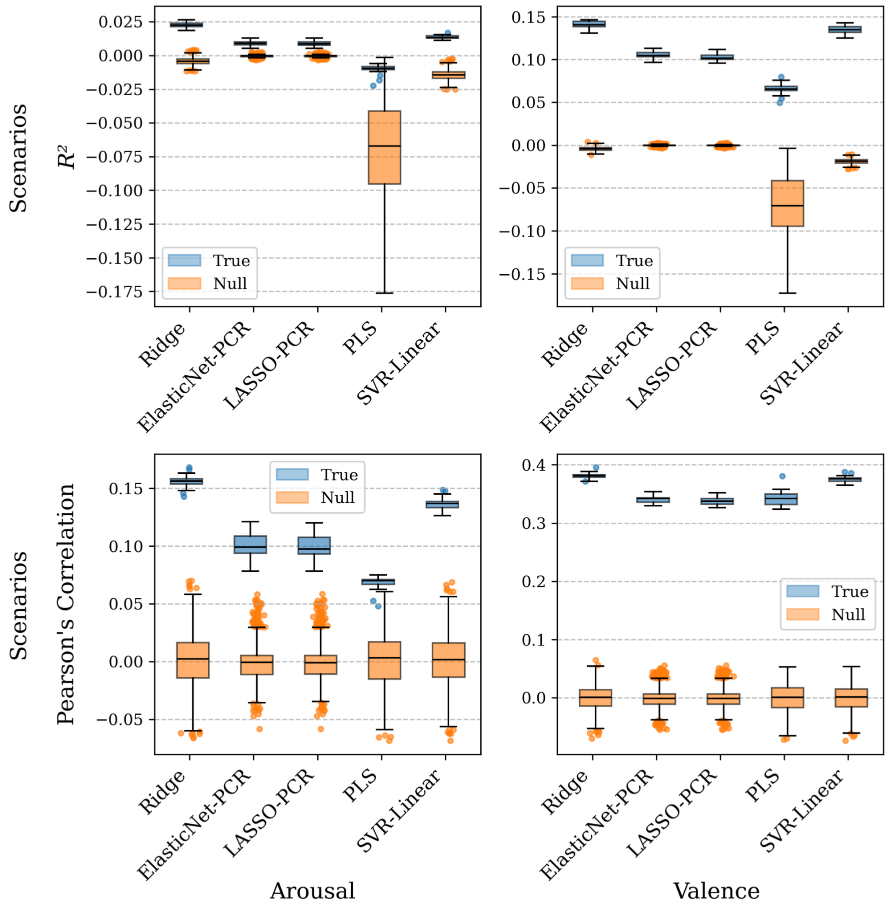


**Fig S7. Boxplots depicting all models' performance in decoding arousal (left) and valence (right) from fMRI data collected during scenario viewing.**

The top row shows $R^2$ and the bottom row shows Pearson's correlation. The distribution of each models' 25 repetitions are shown in blue and their associated 1000 permutations, which were used as a baseline for calculating *p*-value, in orange. The permutation tests indicate that all are significant with a *p*-value of less than 0.001, except decoding of arousal from scenarios using PLS which was significant at a *p*-value of 0.026.

### Example scenario decoding predictions

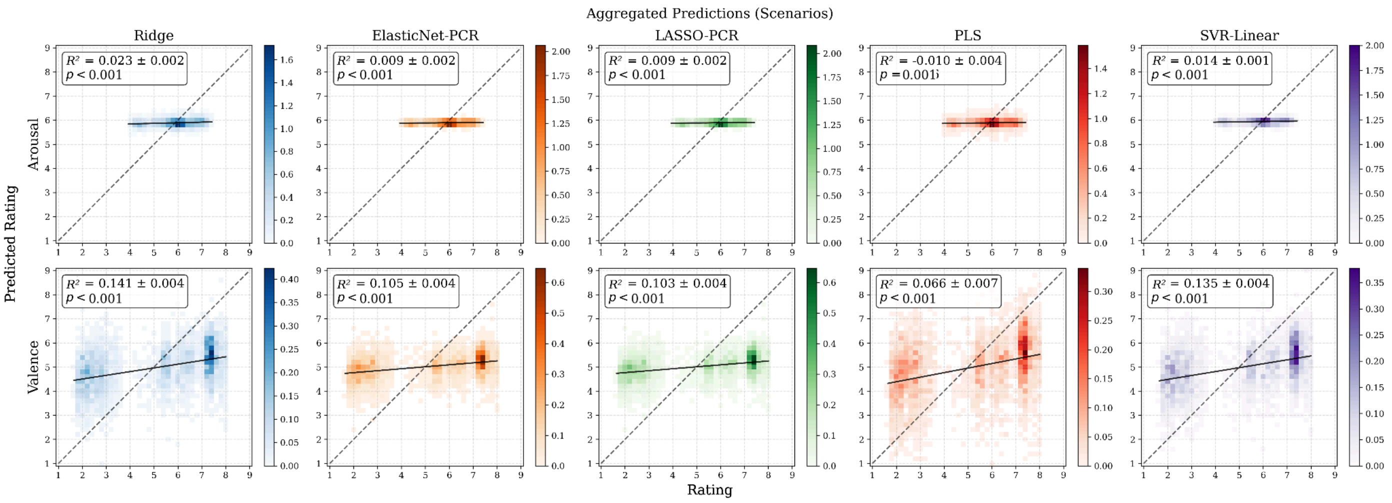


**Fig S8. 2D histograms of the predicted versus rated arousal and valence values for each movie prediction model across all five test sets of a sample five-fold cross-validation.**

The included $R^2$ values were averaged across 25 repetitions of each model. The *p*-values were estimated empirically against 1000 repetitions of permutation tests, making a *p*-value of approximately 0.001 the smallest achievable. The x-axis represents the rated score, and the y-axis represents the model's predicted value. Colorbars are defined using density, which represents the ratio of values present in a bin, divided by the bin size, which is 0.25 in this case. The dotted line indicates a perfect one-to-one correlation.

### FMRIPrep boilerplate

The following boilerplate text was automatically generated by fMRIPrep with the express intention that users should directly copy it into their manuscripts unchanged. It is released under the CC0 license.

Results included in this manuscript come from preprocessing performed with fMRIPrep 23.0.2 (8,9), which is based on Nipype 1.8.6 (10,11).

#### Preprocessing of $B_0$ inhomegeity mappings

A total of 9 fieldmaps were found available within the input BIDS structure for this particular subject. A $B_0$-nonuniformity map (or fieldmap) was estimated based on two (or more) echo-planar imaging (EPI) references with topup ((12); FSL 6.0.5.1).

#### Anatomical data preprocessing

A total of 1 T1-weighted (T1w) images were found within the input BIDS dataset. The T1-weighted (T1w) image was corrected for intensity non-uniformity (INU) with N4BiasFieldCorrection (13), distributed with ANTs 2.3.3 (14), and used as T1w-reference throughout the workflow. The T1w-reference was then skull-stripped with a *Nipype* implementation of the antsBrainExtraction.sh workflow (from ANTs), using MNI152NLin6Asym as target template. Brain tissue segmentation of cerebrospinal fluid (CSF), white-matter (WM) and gray-matter (GM) was performed on the brain-extracted T1w using fast (FSL 6.0.5.1, (15)). Brain surfaces were reconstructed using recon-all (FreeSurfer 7.3.2, (16)), and the brain mask estimated previously was refined with a custom variation of the method to reconcile ANTs-derived and FreeSurfer-derived segmentations of the cortical gray-matter of Mindboggle (17). Volume-based spatial normalization to two standard spaces (MNI152NLin6Asym, MNI152NLin2009cAsym) was performed through nonlinear registration with antsRegistration (ANTs 2.3.3), using brain-extracted versions of both T1w reference and the T1w template. The following templates were were selected for spatial normalization and accessed with *TemplateFlow* (23.0.0, (18)): *FSL's MNI ICBM 152 non-linear 6th Generation Asymmetric Average Brain Stereotaxic Registration Model* (19); TemplateFlow ID: MNI152NLin6Asym], *ICBM 152 Nonlinear Asymmetrical template version 2009c* (20); TemplateFlow ID: MNI152NLin2009cAsym].

#### Functional data preprocessing

For each of the 9 BOLD runs found per subject (across all tasks and sessions), the following preprocessing was performed. First, a reference volume and its skull-stripped version were generated using a custom methodology of *fMRIPrep*. Head-motion parameters with respect to the BOLD reference (transformation matrices, and six corresponding rotation and translation parameters) are estimated before any spatiotemporal filtering using mcflirt (FSL 6.0.5.1:57b01774, (21)). The estimated *fieldmap* was then aligned with rigid-registration to the target EPI (echo-planar imaging) reference run. The field coefficients were mapped on to the reference EPI using the transform. BOLD runs were slice-time corrected to 0.948s (0.5 of slice acquisition range 0s-1.9s) using 3dTshift from AFNI ((22), RRID:SCR_005927). The BOLD reference was then co-registered to the T1w reference using bbregister (FreeSurfer) which implements boundary-based registration (23). Co-registration was configured with six degrees of freedom. Several confounding time-series were calculated based on the *preprocessed BOLD*: framewise displacement (FD), DVARS and three region-wise global signals. FD was computed using two formulations following Power (absolute sum of relative motions, (24)) (relative root mean square displacement between affines, (21)). FD and DVARS are calculated for each functional run, both using their implementations in *Nipype* (following the definitions by (24)). The three global signals are extracted within the CSF, the WM, and the whole-brain masks. Additionally, a set of physiological regressors were extracted to allow for component-based noise correction (*CompCor*, (25)). Principal components are estimated after high-pass filtering the preprocessed BOLD time-series (using a discrete cosine filter with 128s cut-off) for the two CompCor variants: temporal (tCompCor) and anatomical (aCompCor). tCompCor components are then calculated from the top 2% variable voxels within the brain mask. For aCompCor, three probabilistic masks (CSF, WM and combined CSF+WM) are generated in anatomical space. The implementation differs from that of Behzadi et al. in that instead of eroding the masks by 2 pixels on BOLD space, a mask of pixels that likely contain a volume fraction of GM is subtracted from the aCompCor masks. This mask is obtained by dilating a GM mask extracted from the FreeSurfer's aseg segmentation, and it ensures components are not extracted from voxels containing a minimal fraction of GM. Finally, these masks are resampled into BOLD space and binarized by thresholding at 0.99 (as in the original implementation). Components are also calculated separately within the WM and CSF masks. For each CompCor decomposition, the *k* components with the largest singular values are retained, such that the retained components' time series are sufficient to explain 50 percent of variance across the nuisance mask (CSF, WM, combined, or temporal). The remaining components are dropped from consideration. The head-motion estimates calculated in the correction step were also placed within the corresponding confounds file. The confound time series derived from head motion estimates and global signals were expanded with the inclusion of temporal derivatives and quadratic terms for each (26). Frames that exceeded a threshold of 0.5 mm FD or 1.5 standardized DVARS were annotated as motion outliers. Additional nuisance timeseries are calculated by means of principal components analysis of the signal found within a thin band (*crown*) of voxels around the edge of the brain, as proposed by (27). The BOLD time-series were resampled into standard space, generating a *preprocessed BOLD run in MNI152NLin6Asym space*. First, a reference volume and its skull-stripped version were generated using a custom methodology of *fMRIPrep*. Automatic removal of motion artifacts using independent component analysis (ICA-AROMA, (28)) was performed on the *preprocessed BOLD on MNI space* time-series after removal of non-steady state volumes and spatial smoothing with an isotropic, Gaussian kernel of 6mm FWHM (full-width half-maximum). Corresponding "non-aggressively" denoised runs were produced after such smoothing. Additionally, the "aggressive" noise-regressors were collected and placed in the corresponding confounds file. All resamplings can be performed with a *single interpolation step* by composing all the pertinent transformations (i.e. head-motion transform matrices, susceptibility distortion correction when available, and co-registrations to anatomical and output spaces). Gridded (volumetric) resamplings were performed using antsApplyTransforms (ANTs), configured with Lanczos interpolation to minimize the smoothing effects of other kernels (29). Non-gridded (surface) resamplings were performed using mri_vol2surf (FreeSurfer).

#### Full Ridge visualizations

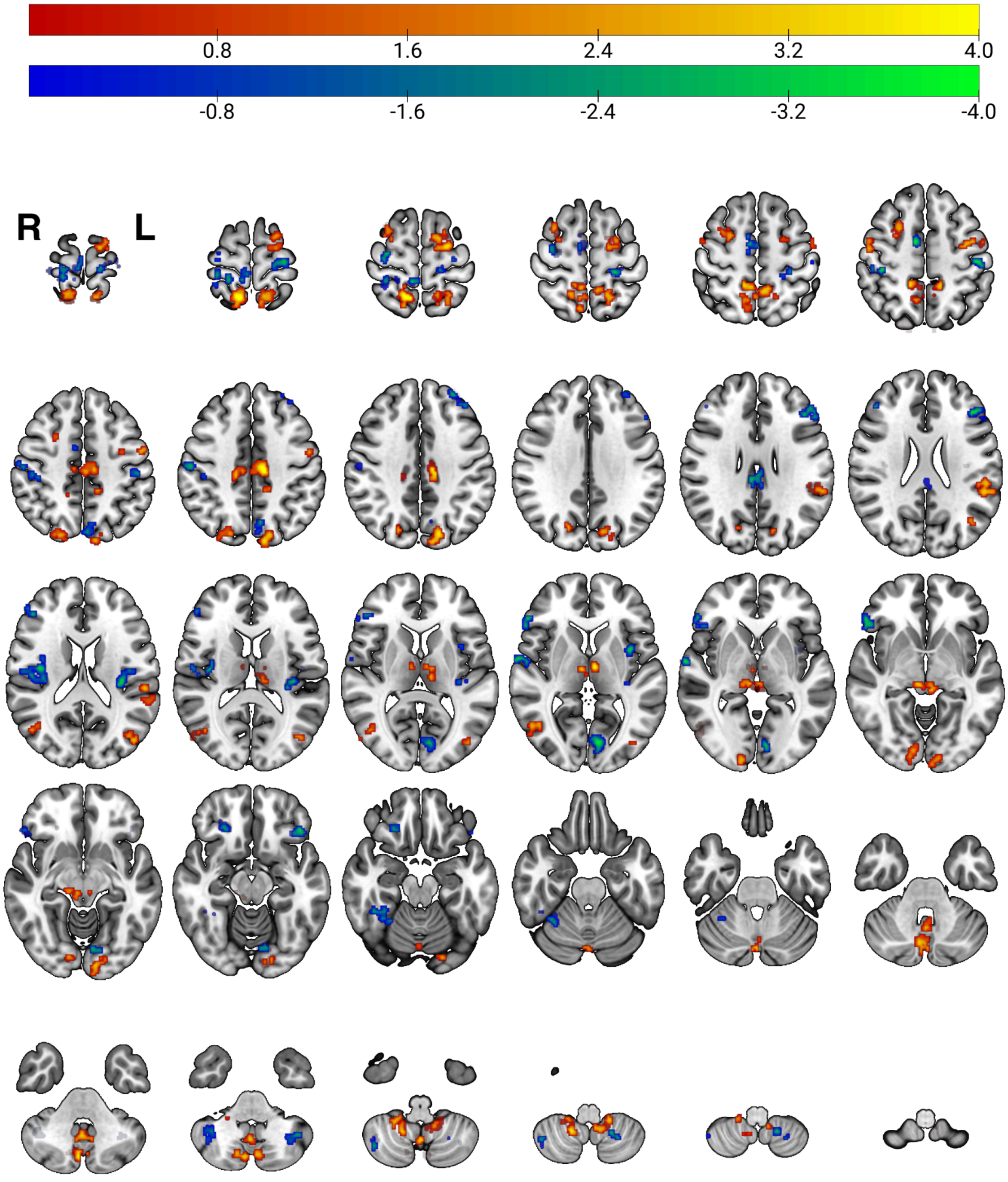


**Fig S9. Evenly spaced axial slices showing key voxels from the Ridge regression model used to decode arousal from fMRI data acquired during movie viewing.**

Coefficients are thresholded at $|z| > 1.96$, and only clusters of ≥30 contiguous voxels (NN=1) are shown. Negative coefficients are displayed in blue-green, positive in red-yellow. The colorbar maximums are set to a z-score of four.

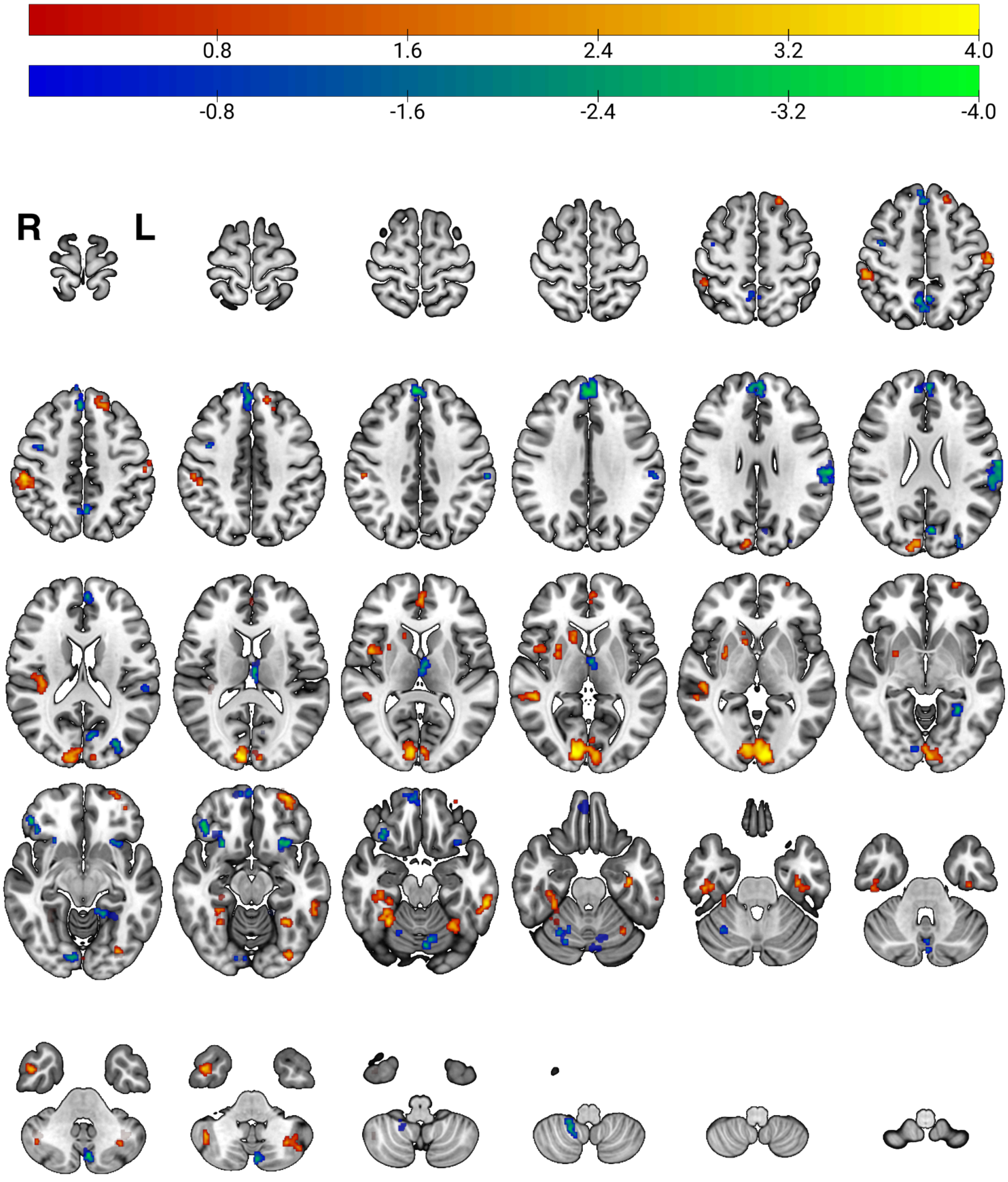


**Fig S10. Evenly spaced axial slices showing key voxels from the Ridge regression model used to decode valence from fMRI data acquired during movie viewing.**

Coefficients are thresholded at $|z| > 1.96$, and only clusters of ≥30 contiguous voxels (NN=1) are shown. Negative coefficients are displayed in blue-green, positive in red-yellow. The colorbar maximums are set to a z-score of four.

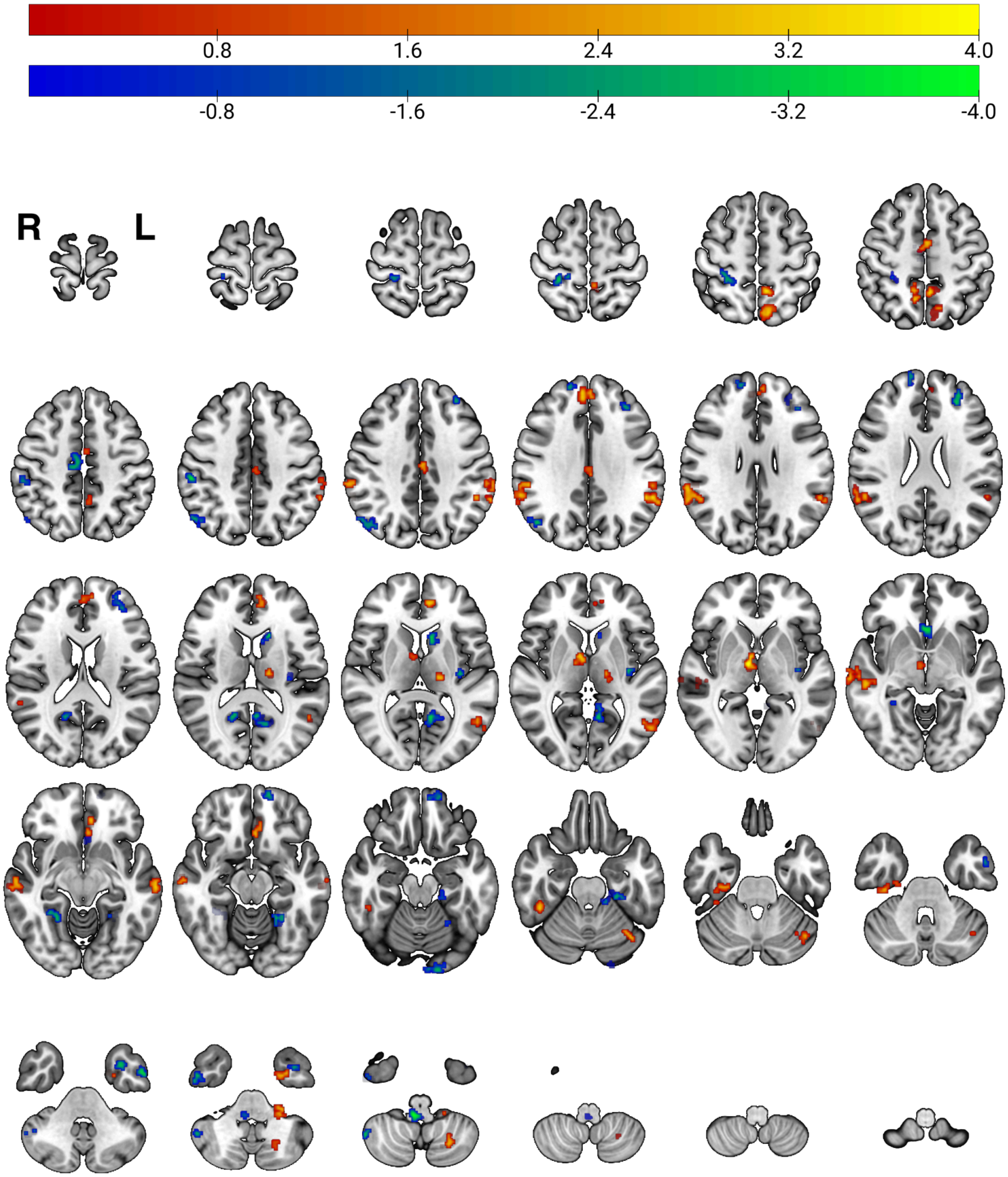


**Fig S11. Evenly spaced axial slices showing key voxels from the Ridge regression model used to decode arousal from fMRI data acquired during scenario viewing.**

Coefficients are thresholded at $|z| > 1.96$, and only clusters of ≥30 contiguous voxels (NN=1) are shown. Negative coefficients are displayed in blue-green, positive in red-yellow. The colorbar maximums are set to a z-score of four. Note that although significant, decoding performance was only slightly better than chance. Refer to the main manuscript for more information.

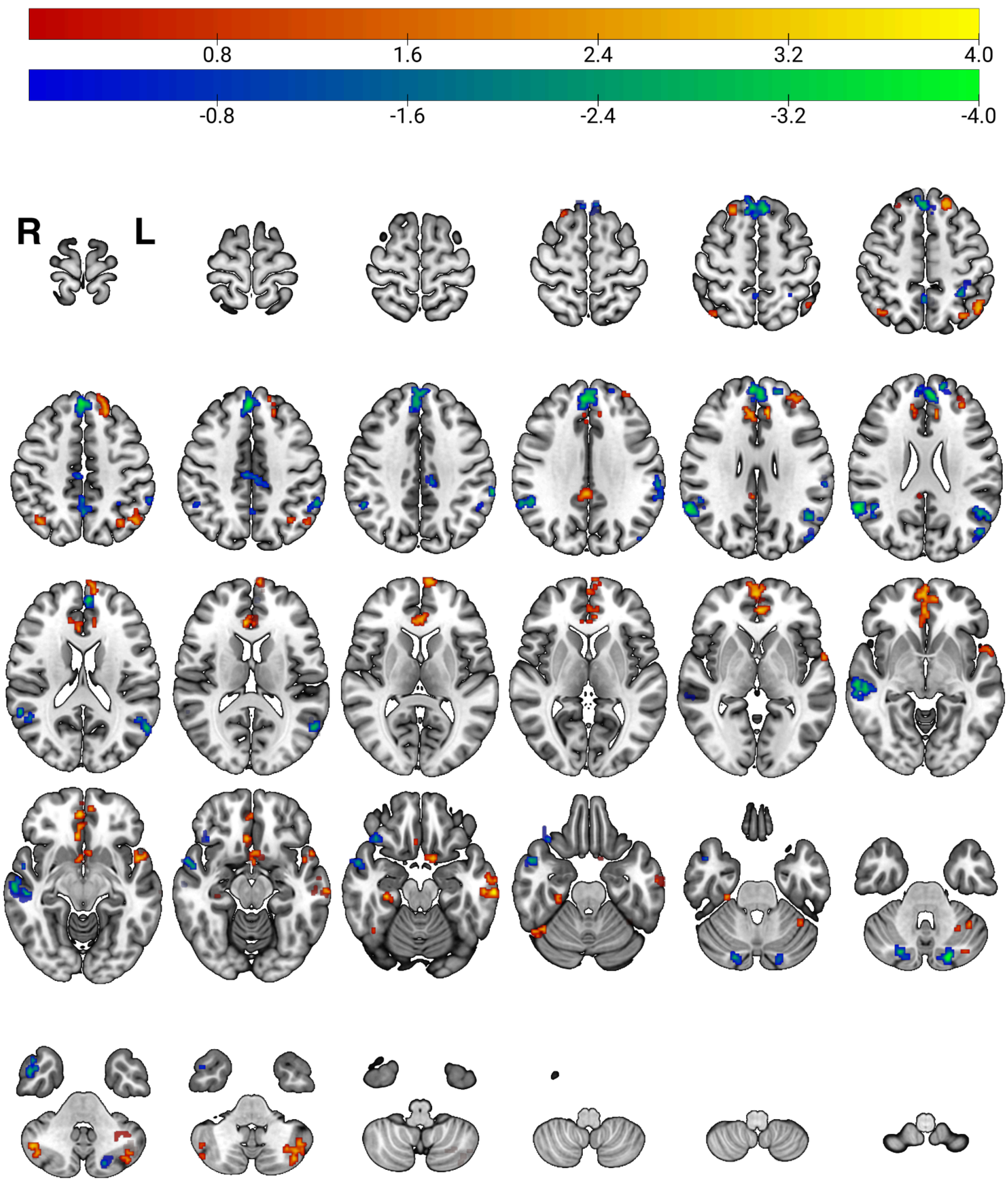


**Fig S12. Evenly spaced axial slices showing key voxels from the Ridge regression model used to decode valence from fMRI data acquired during scenario viewing.**

Coefficients are thresholded at |z| > 1.96, and only clusters of ≥30 contiguous voxels (NN=1) are shown. Negative coefficients are displayed in blue-green, positive in red-yellow. The colorbar maximums are set to a z-score of four.

#### Cluster tables

The following tables (S2-S9) display the cluster local maxima from different modeling (Ridge or parametric modulation), targets (arousal or valence), and induction modality (movies or scenarios). The maxima of each cluster is reported along with cluster size (voxels), peak coordinates (MNI space), hemisphere designation, and corresponding anatomical labels.

**Table S2. Significant Clusters for Voxels from the Ridge Model (Arousal, Movies)**

| Cluster Size (Voxels) | Peak Z | MNI | | | Hemisphere | Anatomical Label |
|---|---|---|---|---|---|---|
| | | x | y | z | | |
| 211 | 5.8 | 12 | -54 | 70 | R | Superior Parietal Lobule |
| 180 | 3.91 | -24 | -8 | 66 | L | Precentral Gyrus |
| 157 | 3.91 | -16 | -80 | 40 | L | Lateral Occipital Cortex, superior division |
| 148 | 4.11 | 4 | -76 | -38 | R | Right Crus II |
| 148 | 3.07 | -12 | -54 | 76 | L | Superior Parietal Lobule |
| 136 | 4.86 | -12 | -22 | 42 | L | Precentral Gyrus |
| 134 | 3.6 | -56 | -30 | 20 | L | Parietal Operculum Cortex |
| 120 | -3.5 | 40 | -12 | 20 | R | Central Opercular Cortex |
| 109 | 3.24 | 24 | 8 | 50 | R | Superior Frontal Gyrus |
| 109 | -3.9 | -8 | -80 | 6 | L | Intracalcarine Cortex |
| 108 | 4.74 | 0 | -60 | -46 | C | Vermis IX |
| 108 | -3.45 | 54 | 32 | -6 | R | Inferior Frontal Gyrus, pars triangularis |
| 106 | 3.89 | 6 | -28 | -2 | R | Brain-Stem |
| 105 | 4.35 | 16 | -54 | -50 | R | Right VIIIb |
| 94 | 3.32 | -14 | -42 | 52 | L | Postcentral Gyrus |
| 94 | 2.99 | 52 | -64 | 4 | R | Lateral Occipital Cortex, inferior division |
| 92 | 3.23 | -12 | -92 | -10 | L | Occipital Pole |
| 86 | -4.02 | -36 | -24 | 16 | L | Parietal Operculum Cortex |
| 85 | 3.44 | 20 | -74 | 36 | R | Precuneus Cortex |
| 82 | 4.32 | 14 | -42 | 54 | R | Postcentral Gyrus |
| 78 | -2.71 | 42 | -44 | -18 | R | Temporal Occipital Fusiform Cortex |
| 78 | 3.16 | -46 | -78 | 20 | L | Lateral Occipital Cortex, superior division |
| 71 | -4 | 10 | -4 | 52 | R | Juxtapositional Lobule Cortex (formerly Supplementary Motor Cortex) |
| 70 | -3.17 | -30 | -22 | 72 | L | Precentral Gyrus |
| 68 | -3.93 | -46 | 32 | 26 | L | Middle Frontal Gyrus |
| 64 | 3.41 | 14 | -94 | -2 | R | Occipital Pole |
| 63 | 3.37 | -20 | -44 | -52 | L | Left VIIIb |
| 59 | -2.85 | 16 | -30 | 78 | R | Postcentral Gyrus |

| | | | | | | |
|---|---|---|---|---|---|---|
| 54 | -3.75 | 64 | -4 | 4 | R | Planum Polare |
| 54 | -3.65 | 8 | -40 | 68 | R | Postcentral Gyrus |
| 54 | 3.51 | 8 | -64 | 60 | R | Precuneus Cortex |
| 49 | -4.06 | -46 | -22 | 52 | L | Postcentral Gyrus |
| 48 | -2.86 | 46 | 36 | 20 | R | Frontal Pole |
| 45 | -3.23 | 32 | -20 | 68 | R | Precentral Gyrus |
| 44 | -3.54 | 6 | -30 | 28 | R | Cingulate Gyrus, posterior division |
| 44 | -2.89 | -8 | -78 | -10 | L | Lingual Gyrus |
| 43 | -3.49 | 40 | -30 | 52 | R | Postcentral Gyrus |
| 42 | -3.24 | 26 | 34 | -16 | R | Frontal Pole |
| 42 | -3.14 | -30 | 48 | 38 | L | Frontal Pole |
| 40 | -3.01 | 56 | -18 | 42 | R | Postcentral Gyrus |
| 39 | -2.9 | 32 | -40 | 68 | R | Superior Parietal Lobule |
| 37 | 3.39 | -52 | -6 | 44 | L | Precentral Gyrus |
| 37 | -3.15 | -8 | -68 | 44 | L | Precuneus Cortex |
| 36 | 4.39 | -6 | -10 | 6 | L | Left Thalamus |
| 35 | 3.12 | -42 | -4 | 52 | L | Precentral Gyrus |
| 35 | -3.53 | -18 | -54 | -56 | L | Left VIIIb |
| 34 | -3.43 | -24 | -32 | 62 | L | Postcentral Gyrus |
| 33 | 4.09 | 16 | -26 | 42 | R | Precentral Gyrus |
| 33 | 3.17 | 6 | -12 | 6 | R | Right Thalamus |
| 33 | -2.78 | 42 | -66 | -48 | R | Right Crus II |
| 33 | -2.69 | -36 | -62 | -44 | L | Left Crus II |
| 32 | -3.98 | -38 | 4 | 4 | L | Insular Cortex |
| 32 | -3.72 | -44 | 30 | -14 | L | Frontal Orbital Cortex |
| 31 | 3.01 | -6 | -74 | -44 | L | Left VIIb |
| 30 | 3.3 | -10 | -20 | 10 | L | Left Thalamus |
| 30 | 2.77 | 50 | -2 | 54 | R | Precentral Gyrus |
| 30 | -3.14 | 36 | -52 | -42 | R | Right Crus II |

| Table S3. Significant Clusters for Voxels from the Ridge Model (Valence, Movies) | | | | | | |
|---|---|---|---|---|---|---|
| Cluster Size (Voxels) | Peak Z | MNI | | | Hemisphere | Anatomical Label |
| | | x | y | z | | |
| 709 | 5.55 | 14 | -86 | 4 | R | Intracalcarine Cortex |
| 376 | -3.83 | 4 | 50 | 38 | R | Superior Frontal Gyrus |
| 189 | -3.61 | -66 | -24 | 26 | L | Supramarginal Gyrus, anterior division |
| 130 | 3.85 | 54 | -28 | 48 | R | Supramarginal Gyrus, anterior division |
| 90 | -3.3 | -4 | -56 | 50 | L | Precuneus Cortex |
| 85 | 3.96 | 28 | -52 | -20 | R | Right VI |
| 78 | 4.31 | -32 | 60 | -14 | L | Frontal Pole |
| 77 | -4.04 | -2 | -16 | 12 | L | Left Thalamus |
| 76 | 3.45 | 50 | -38 | 4 | R | Middle Temporal Gyrus, temporooccipital part |
| 72 | 4.5 | 36 | -28 | -22 | R | Temporal Fusiform Cortex, posterior division |
| 63 | 4.77 | -60 | -36 | -18 | L | Inferior Temporal Gyrus, posterior division |
| 63 | 3.59 | -32 | -52 | -16 | L | Temporal Occipital Fusiform Cortex |
| 61 | -3.14 | 6 | 50 | -22 | R | Frontal Medial Cortex |
| 60 | -3.82 | 44 | 36 | -12 | R | Frontal Pole |
| 60 | -3.66 | -4 | -76 | -40 | L | Left Crus II / Left VIIb |
| 59 | -3.81 | -26 | 20 | -14 | L | Frontal Orbital Cortex |
| 54 | 3.05 | -20 | 34 | 56 | L | Superior Frontal Gyrus |
| 54 | 3.6 | 44 | 2 | -42 | R | Inferior Temporal Gyrus, anterior division |
| 51 | -3.6 | 26 | 18 | -14 | R | Frontal Orbital Cortex |
| 49 | -3.59 | -4 | -74 | -20 | L | Vermis VI |
| 48 | -3.33 | -28 | -52 | -4 | L | Lingual Gyrus |
| 46 | 3.68 | -30 | -64 | -42 | L | Left Crus II |
| 45 | 3.08 | 40 | -22 | 20 | R | Parietal Operculum Cortex |
| 44 | -2.94 | -30 | -80 | 18 | L | Lateral Occipital Cortex, superior division |
| 42 | -2.7 | 20 | -62 | -22 | R | Right VI |
| 42 | -3.02 | 14 | -86 | -12 | R | Occipital Fusiform Gyrus |
| 42 | 3.93 | -36 | -16 | -26 | L | Temporal Fusiform Cortex, posterior division |
| 41 | 2.9 | 28 | 2 | -2 | R | Right Putamen |
| 39 | -4.14 | -6 | -76 | 22 | L | Cuneal Cortex |
| 38 | 3.2 | 44 | 4 | 8 | R | Central Opercular Cortex |

| | | | | | | |
|---|---|---|---|---|---|---|
| 37 | 3.48 | 42 | -56 | -40 | R | Right Crus I |
| 36 | -3.25 | 16 | -52 | -52 | R | Right VIIIb |
| 35 | 3.34 | 48 | -22 | -28 | R | Inferior Temporal Gyrus, posterior division |
| 33 | 3.08 | 0 | 48 | 10 | C | Paracingulate Gyrus |
| 33 | 2.87 | -58 | -20 | 52 | L | Postcentral Gyrus |
| 31 | 3.07 | 14 | 14 | 4 | R | Right Caudate |
| 30 | 3.17 | -32 | -82 | -12 | L | Occipital Fusiform Gyrus |
| 30 | -2.75 | 40 | -6 | 54 | R | Precentral Gyrus |

**Table S4. Significant Clusters for Voxels from the Ridge Model (Arousal, Scenarios)**

| Cluster Size (Voxels) | Peak Z | MNI | | | Hemisphere | Anatomical Label |
|---|---|---|---|---|---|---|
| | | x | y | z | | |
| 194 | 3.48 | 62 | -48 | 26 | R | Angular Gyrus |
| 101 | -3.67 | -10 | -58 | 8 | L | Precuneus Cortex |
| 99 | 4.26 | -52 | -46 | 36 | L | Supramarginal Gyrus, posterior division |
| 90 | 5.03 | 8 | -8 | 0 | R | Right Thalamus |
| 87 | -3.31 | 52 | -66 | 44 | R | Lateral Occipital Cortex, superior division |
| 83 | -3.22 | 26 | -38 | 64 | R | Postcentral Gyrus |
| 82 | 3.5 | 62 | -16 | -12 | R | Middle Temporal Gyrus, posterior division |
| 70 | 3.57 | -10 | -68 | 56 | L | Lateral Occipital Cortex, superior division |
| 67 | 3.81 | 6 | 48 | 32 | R | Superior Frontal Gyrus |
| 66 | 3.47 | -8 | 46 | 10 | L | Paracingulate Gyrus |
| 66 | 4.01 | -8 | -48 | 56 | L | Precuneus Cortex |
| 64 | -3.48 | -28 | 40 | 24 | L | Frontal Pole |
| 55 | -3.07 | 14 | 54 | 36 | R | Frontal Pole |
| 52 | 2.8 | -58 | -66 | 6 | L | Lateral Occipital Cortex, inferior division |
| 51 | -3.26 | 6 | -18 | 48 | R | Precentral Gyrus |
| 48 | 3.29 | -66 | -36 | 40 | L | Supramarginal Gyrus, anterior division |
| 47 | 4.18 | -4 | 26 | -12 | L | Subcallosal Cortex |
| 41 | 3.07 | -2 | -20 | 36 | L | Cingulate Gyrus, posterior division |
| 41 | -3.6 | -16 | 60 | -18 | L | Frontal Pole |
| 40 | -5.01 | 6 | -40 | -46 | R | Brain-Stem |
| 39 | -4.32 | -16 | 18 | 14 | L | Left Caudate |
| 38 | -3.08 | -22 | -50 | -14 | L | Temporal Occipital Fusiform Cortex |
| 38 | -3.63 | 0 | 22 | -6 | C | Subcallosal Cortex |
| 38 | 3.28 | -40 | -68 | -26 | L | Left Crus I |

| | | | | | | |
|---|---|---|---|---|---|---|
| 37 | -3.29 | 24 | -50 | -10 | R | Lingual Gyrus |
| 37 | 3.43 | -16 | -16 | 12 | L | Left Thalamus |
| 36 | -3 | -26 | -30 | -24 | L | Parahippocampal Gyrus, posterior division |
| 36 | 4.41 | 38 | -26 | -32 | R | Temporal Fusiform Cortex, posterior division |
| 35 | -3.64 | 50 | -56 | -46 | R | Right Crus II |
| 35 | -3.19 | -18 | -98 | -20 | L | Occipital Pole |
| 34 | -3.5 | -36 | -16 | 8 | L | Insular Cortex |
| 34 | -3.65 | 46 | -6 | -46 | R | Inferior Temporal Gyrus, anterior division |
| 34 | 3.3 | -24 | -40 | -44 | L | Left X |
| 34 | 3.04 | 54 | -26 | -6 | R | Middle Temporal Gyrus, posterior division |
| 34 | 3.43 | 10 | -56 | 52 | R | Precuneus Cortex |
| 34 | -4.21 | -54 | 0 | -36 | L | Middle Temporal Gyrus, anterior division |
| 33 | -3.54 | -34 | 6 | -40 | L | Temporal Pole |
| 33 | -3.58 | 14 | -54 | 16 | R | Precuneus Cortex |
| 33 | 3.64 | -28 | -4 | -42 | L | Temporal Fusiform Cortex, anterior division |
| 33 | 3.34 | 46 | -40 | -22 | R | Inferior Temporal Gyrus, temporooccipital part |
| 32 | 3.51 | -66 | -22 | -10 | L | Middle Temporal Gyrus, posterior division |
| 32 | 3.73 | -26 | -66 | -46 | L | Left VIIb |
| 31 | 3.17 | -2 | -4 | 52 | L | Juxtapositional Lobule Cortex (formerly Supplementary Motor Cortex) |
| 30 | -4.48 | 54 | -30 | 44 | R | Supramarginal Gyrus, anterior division |
| 30 | -3.14 | -36 | 34 | 30 | L | Middle Frontal Gyrus |

**Table S5. Significant Clusters for Voxels from the Ridge Model (Valence, Scenarios)**

| Cluster Size (Voxels) | Peak Z | MNI | | | Hemisphere | Anatomical Label |
|---|---|---|---|---|---|---|
| | | x | y | z | | |
| 701 | -4.07 | 2 | 42 | 42 | R | Superior Frontal Gyrus |
| 313 | 3.62 | 2 | 60 | -2 | R | Frontal Pole |
| 228 | -4.58 | 62 | -52 | 24 | R | Angular Gyrus |
| 162 | -4.39 | 62 | -20 | -8 | R | Middle Temporal Gyrus, posterior division |
| 149 | -3.74 | -56 | -62 | 16 | L | Lateral Occipital Cortex, superior division |
| 124 | 3.49 | -32 | -66 | -44 | L | Left Crus II |
| 92 | 3.3 | -20 | 38 | 50 | L | Superior Frontal Gyrus |
| 86 | 3.64 | 6 | 32 | 12 | R | Cingulate Gyrus, anterior division |
| 82 | 3.32 | -48 | -62 | 52 | L | Lateral Occipital Cortex, superior division |

| | | | | | | |
|---|---|---|---|---|---|---|
| 80 | -3.49 | 58 | 6 | -16 | R | Temporal Pole |
| 76 | 3.74 | -10 | 60 | 18 | L | Frontal Pole |
| 71 | -2.86 | 2 | -52 | 50 | R | Precuneus Cortex |
| 68 | -4.8 | -20 | -82 | -34 | L | Left Crus II |
| 67 | 3.46 | -56 | 10 | -8 | L | Temporal Pole |
| 66 | -3.38 | -66 | -36 | 38 | L | Supramarginal Gyrus, anterior division |
| 62 | 3.11 | -8 | 8 | -18 | L | Subcallosal Cortex |
| 58 | 4.81 | -66 | -22 | -18 | L | Middle Temporal Gyrus, posterior division |
| 54 | 2.89 | 6 | 32 | 28 | R | Paracingulate Gyrus |
| 51 | -2.95 | 4 | -22 | 44 | R | Cingulate Gyrus, posterior division |
| 45 | -3.16 | 48 | 2 | -36 | R | Inferior Temporal Gyrus, anterior division |
| 44 | 3.18 | 40 | -64 | 48 | R | Lateral Occipital Cortex, superior division |
| 42 | -4.01 | 22 | -76 | -32 | R | Right Crus I |
| 41 | -3.34 | 38 | 26 | -22 | R | Frontal Orbital Cortex |
| 39 | 3.91 | 46 | -64 | -40 | R | Right Crus I |
| 37 | 3.91 | 30 | -26 | -18 | R | Parahippocampal Gyrus, posterior division |
| 36 | -3.01 | -58 | -46 | 44 | L | Supramarginal Gyrus, posterior division |
| 35 | -2.98 | 48 | -54 | 22 | R | Angular Gyrus |
| 35 | 3.71 | 6 | -42 | 32 | R | Cingulate Gyrus, posterior division |
| 35 | 2.88 | -34 | 52 | 30 | L | Frontal Pole |
| 34 | 2.92 | -40 | -54 | -36 | L | Left Crus I |
| 33 | 3.86 | -68 | -12 | -20 | L | Middle Temporal Gyrus, posterior division |
| 32 | 3.11 | -34 | -68 | 50 | L | Lateral Occipital Cortex, superior division |
| 32 | 3.82 | -10 | 34 | 28 | L | Paracingulate Gyrus |
| 32 | 3.5 | 42 | -56 | -22 | R | Temporal Occipital Fusiform Cortex |
| 31 | -2.96 | -30 | -48 | 54 | L | Superior Parietal Lobule |
| 31 | -2.9 | -50 | -74 | 24 | L | Lateral Occipital Cortex, superior division |
| 31 | -3.24 | -16 | 58 | 24 | L | Frontal Pole |
| 30 | 3.32 | 22 | 28 | 58 | R | Superior Frontal Gyrus |

**Table S6. Significant Clusters for Voxels from the Parametric Modulation (Arousal, Movies)**

| Cluster Size (Voxels) | Peak Z | MNI | | | Hemisphere | Anatomical Label |
|---|---|---|---|---|---|---|
| | | x | y | z | | |
| 32319 | 11.9 | -14 | -24 | 40 | L | Precentral Gyrus |
| 5886 | 10.4 | -26 | -4 | 60 | L | Superior Frontal Gyrus |
| 2570 | -7.15 | -22 | -48 | 20 | L | Left Cerebral White Matter |
| 1659 | -7.55 | -2 | -32 | 62 | L | Precentral Gyrus |

| | | | | | | |
|---|---|---|---|---|---|---|
| 933 | -8.02 | 24 | -100 | -4 | R | Occipital Pole |
| 894 | -7.16 | -38 | -66 | 50 | L | Lateral Occipital Cortex, superior division |
| 883 | -6.02 | 38 | -78 | -42 | R | Right Crus II |
| 810 | -6.83 | 38 | -16 | 18 | R | Central Opercular Cortex |
| 496 | -7.51 | 48 | -60 | 48 | R | Lateral Occipital Cortex, superior division |
| 494 | -6.27 | -36 | -16 | 20 | L | Central Opercular Cortex |
| 455 | 5.77 | 52 | -26 | -6 | R | Middle Temporal Gyrus, posterior division |
| 396 | -5.26 | -42 | 46 | -10 | L | Frontal Pole |
| 325 | -5.63 | -40 | -74 | -46 | L | Left Crus II |
| 233 | 5.68 | 6 | 46 | 38 | R | Superior Frontal Gyrus |
| 188 | -5.22 | -64 | -34 | 2 | L | Superior Temporal Gyrus, posterior division |
| 182 | -7.74 | -22 | -102 | -6 | L | Occipital Pole |
| 138 | -5.65 | -22 | 48 | -12 | L | Frontal Pole |
| 136 | -5.13 | -22 | 34 | 52 | L | Superior Frontal Gyrus |
| 130 | -5.83 | 20 | 40 | -16 | R | Frontal Pole |
| 102 | -4.8 | -4 | 22 | 44 | L | Paracingulate Gyrus |
| 100 | 5.82 | 28 | 16 | -16 | R | Frontal Orbital Cortex |
| 97 | -4.38 | -12 | 56 | 36 | L | Frontal Pole |
| 87 | 4.86 | -34 | 34 | 44 | L | Middle Frontal Gyrus |
| 58 | -4.74 | 46 | 34 | 32 | R | Middle Frontal Gyrus |
| 46 | 4.27 | 6 | 22 | 62 | R | Superior Frontal Gyrus |
| 45 | -4.57 | 18 | 30 | 0 | R | Right Cerebral White Matter |
| 44 | -4.6 | 56 | -36 | -14 | R | Inferior Temporal Gyrus, posterior division |
| 42 | -3.99 | 42 | 50 | 8 | R | Frontal Pole |
| 41 | 4.88 | -32 | 18 | 12 | L | Frontal Operculum Cortex |
| 38 | -4.41 | 22 | 58 | 0 | R | Frontal Pole |
| 35 | 4.42 | 22 | 14 | 0 | R | Right Putamen |
| 35 | -4.98 | -26 | 22 | 60 | L | Superior Frontal Gyrus |
| 35 | -5.91 | -12 | -66 | -4 | L | Lingual Gyrus |
| 32 | 5.74 | 20 | -2 | -12 | R | Right Amygdala |
| 32 | -5.56 | 18 | 24 | 62 | R | Superior Frontal Gyrus |
| 31 | -4.83 | 16 | 4 | 24 | R | Right Caudate |
| 29 | -4.18 | -16 | 26 | 10 | L | Left Lateral Ventrical |
| 29 | 4.41 | -24 | 10 | 4 | L | Left Putamen |
| 29 | -4.59 | -48 | 26 | 30 | L | Middle Frontal Gyrus |

| Table S7. Significant Clusters for Voxels from the Parametric Modulation (Valence, Movies) | | | | | | |
|---|---|---|---|---|---|---|
| **Cluster Size (Voxels)** | **Peak Z** | **MNI** | | | **Hemisphere** | **Anatomical Label** |
| | | **x** | **y** | **z** | | |
| 4422 | -7.77 | 4 | -52 | 52 | R | Precuneus Cortex |
| 3795 | 8.8 | 12 | -92 | 18 | R | Occipital Pole |
| 3553 | 7.43 | -30 | -70 | 46 | L | Lateral Occipital Cortex, superior division |
| 2096 | -7.01 | 56 | -66 | 10 | R | Lateral Occipital Cortex, inferior division |
| 1683 | -7.41 | -8 | -76 | -44 | L | Left VIIb |
| 1551 | -8.36 | 4 | 46 | 40 | R | Superior Frontal Gyrus |
| 1543 | -7.6 | 30 | 20 | -14 | R | Frontal Orbital Cortex |
| 1337 | 7.18 | -42 | 50 | 10 | L | Frontal Pole |
| 840 | -6.34 | -42 | -62 | 16 | L | Lateral Occipital Cortex, inferior division |
| 508 | -6.78 | -64 | -32 | 22 | L | Supramarginal Gyrus, anterior division |
| 426 | -5.7 | 52 | 0 | -18 | R | Superior Temporal Gyrus, anterior division |
| 406 | -5.96 | 2 | 10 | 32 | R | Cingulate Gyrus, anterior division |
| 389 | 5.57 | 28 | -70 | 50 | R | Lateral Occipital Cortex, superior division |
| 357 | -7.93 | -30 | 18 | -14 | L | Frontal Orbital Cortex |
| 232 | -5.99 | 14 | -48 | -52 | R | Right IX |
| 147 | 5.98 | 38 | -14 | 18 | R | Central Opercular Cortex |
| 134 | 5.73 | -54 | -56 | -16 | L | Inferior Temporal Gyrus, temporooccipital part |
| 132 | 5.64 | 38 | -60 | -42 | R | Right Crus II |
| 131 | -4.63 | -34 | -46 | -44 | L | Left VIIb |
| 128 | -5.42 | 2 | 54 | -14 | R | Frontal Medial Cortex |
| 128 | -5.79 | 4 | -32 | -28 | R | Brain-Stem |
| 120 | 5.79 | 50 | -46 | -8 | R | Inferior Temporal Gyrus, temporooccipital part |
| 117 | 5.6 | -24 | 46 | -16 | L | Frontal Pole |
| 116 | 5.26 | 18 | 2 | 26 | R | Right Cerebral White Matter |
| 109 | 5.01 | 36 | 52 | -4 | R | Frontal Pole |
| 99 | 4.92 | -2 | 18 | 2 | L | Subcallosal Cortex |
| 93 | -5.39 | 40 | 0 | -14 | R | Insular Cortex |
| 90 | -4.39 | 28 | -80 | -34 | R | Right Crus I |
| 79 | 6.96 | -2 | -32 | 36 | L | Cingulate Gyrus, posterior division |
| 76 | 4.61 | 20 | 42 | -4 | R | Right Cerebral White Matter |
| 68 | -5.25 | -34 | 22 | 8 | L | Frontal Operculum Cortex |
| 67 | 4.21 | 42 | 40 | 22 | R | Frontal Pole |

| | | | | | | |
|---|---|---|---|---|---|---|
| 61 | -4.88 | 18 | 4 | 70 | R | Superior Frontal Gyrus |
| 59 | -4.14 | -40 | -4 | 50 | L | Precentral Gyrus |
| 54 | -5.27 | 44 | -48 | -22 | R | Temporal Occipital Fusiform Cortex |
| 51 | 4.65 | -38 | -14 | 20 | L | Central Opercular Cortex |
| 50 | -4.8 | -12 | -30 | 38 | L | Cingulate Gyrus, posterior division |
| 47 | -5.05 | 12 | -72 | -18 | R | Right VI |
| 47 | 4.16 | 50 | -18 | 42 | R | Postcentral Gyrus |
| 41 | 5.08 | -18 | 32 | 58 | L | Superior Frontal Gyrus |
| 39 | -4.69 | 18 | -74 | 28 | R | Cuneal Cortex |
| 38 | 4.24 | -38 | -66 | -42 | L | Left Crus II |
| 31 | 3.97 | -20 | -28 | 28 | L | Left Cerebral White Matter |
| 29 | 5.08 | 16 | 64 | 2 | R | Frontal Pole |

**Table S8. Significant Clusters for Voxels from the Parametric Modulation (Arousal, Scenarios)**

| Cluster Size (Voxels) | Peak Z | MNI | | | Hemisphere | Anatomical Label |
|---|---|---|---|---|---|---|
| | | x | y | z | | |
| 3706 | 6.64 | -2 | -90 | -10 | L | Lingual Gyrus |
| 645 | 5.1 | -2 | -56 | 66 | L | Precuneus Cortex |
| 612 | 5.83 | -56 | -66 | 18 | L | Lateral Occipital Cortex, superior division |
| 143 | 4.54 | 42 | -64 | 28 | R | Lateral Occipital Cortex, superior division |
| 142 | 4.57 | -40 | -44 | -22 | L | Temporal Fusiform Cortex, posterior division |
| 110 | 5.31 | 8 | -76 | -40 | R | Right Crus II |
| 91 | 4.69 | 2 | -30 | -2 | R | Brain-Stem |
| 77 | 4.18 | 4 | 48 | 30 | R | Superior Frontal Gyrus |
| 74 | 4.67 | 62 | -46 | 20 | R | Angular Gyrus |
| 49 | 4.27 | 8 | 56 | 10 | R | Frontal Pole |
| 41 | 4.28 | 28 | 32 | 46 | R | Middle Frontal Gyrus |
| 36 | 4.28 | 40 | -38 | -22 | R | Temporal Fusiform Cortex, posterior division |
| 34 | 4.24 | 58 | -26 | -2 | R | Superior Temporal Gyrus, posterior division |
| 30 | 4.47 | -32 | -84 | 24 | L | Lateral Occipital Cortex, superior division |

**Table S9. Significant Clusters for Voxels from the Parametric Modulation (Valence, Scenarios)**

| Cluster Size (Voxels) | Peak Z | MNI | | | Hemisphere | Anatomical Label |
|---|---|---|---|---|---|---|
| | | x | y | z | | |
| 2672 | -8.28 | -50 | -58 | 22 | L | Angular Gyrus |

| | | | | | | |
|---|---|---|---|---|---|---|
| 2456 | -7.93 | -2 | 48 | 34 | L | Superior Frontal Gyrus |
| 2428 | -7.92 | 54 | 6 | -20 | R | Temporal Pole |
| 887 | -6.61 | -54 | 22 | 8 | L | Inferior Frontal Gyrus, pars triangularis |
| 765 | -6.66 | 48 | 34 | -8 | R | Frontal Pole |
| 621 | -7.88 | -20 | -78 | -32 | L | Left Crus I |
| 560 | -6.14 | -2 | -52 | 42 | L | Precuneus Cortex |
| 476 | -6.82 | 20 | -80 | -34 | R | Right Crus II |
| 473 | 5.01 | 48 | -38 | 58 | R | Supramarginal Gyrus, posterior division |
| 354 | 4.9 | 48 | 42 | 22 | R | Frontal Pole |
| 352 | 5.71 | -10 | -64 | 36 | L | Precuneus Cortex |
| 343 | 5.07 | -36 | -56 | 38 | L | Angular Gyrus |
| 330 | 5.08 | 16 | -66 | 40 | R | Precuneus Cortex |
| 318 | 6.09 | 2 | -32 | 28 | R | Cingulate Gyrus, posterior division |
| 305 | -5.5 | 4 | -54 | -40 | R | Right IX |
| 219 | -5.4 | -12 | -2 | 16 | L | Left Caudate |
| 217 | -5.29 | -38 | -2 | 46 | L | Precentral Gyrus |
| 215 | 5.46 | 42 | -64 | -40 | R | Right Crus I |
| 192 | 5.5 | -22 | 30 | -14 | L | Frontal Orbital Cortex |
| 123 | -5.39 | 8 | 10 | 10 | R | Right Caudate |
| 117 | -5.56 | 34 | 6 | 38 | R | Middle Frontal Gyrus |
| 95 | 4.77 | -36 | -68 | -42 | L | Left Crus II |
| 93 | -4.57 | 18 | 62 | 22 | R | Frontal Pole |
| 91 | 4.27 | -10 | 34 | 26 | L | Paracingulate Gyrus |
| 87 | 4.42 | 2 | 16 | 4 | R | Right Lateral Ventricle |
| 82 | 5.21 | -44 | 48 | 14 | L | Frontal Pole |
| 71 | -5.17 | 2 | 56 | -16 | R | Frontal Pole |
| 71 | 4.23 | 28 | 62 | 2 | R | Frontal Pole |
| 63 | 4.65 | 6 | 28 | 18 | R | Cingulate Gyrus, anterior division |
| 58 | 4.36 | 10 | 34 | 24 | R | Paracingulate Gyrus |
| 52 | -4.96 | 0 | -20 | 40 | C | Cingulate Gyrus, posterior division |
| 51 | 4.42 | 32 | 18 | 10 | R | Insular Cortex / Frontal Operculum Cortex |
| 45 | 4.29 | -30 | 38 | 32 | L | Middle Frontal Gyrus |
| 44 | -4.15 | 34 | -92 | 4 | R | Occipital Pole |
| 42 | 4.05 | 66 | -22 | 16 | R | Planum Temporale |
| 41 | 4.74 | -58 | -48 | -14 | L | Inferior Temporal Gyrus, temporooccipital part |
| 39 | -4.88 | 10 | -30 | -2 | R | Right Thalamus |

| | | | | | | |
|---|---|---|---|---|---|---|
| 37 | -4.86 | -36 | 16 | 26 | L | Inferior Frontal Gyrus, pars opercularis |
| 37 | 4.28 | -28 | -36 | 60 | L | Postcentral Gyrus |
| 33 | 4.03 | 16 | -36 | 42 | R | Precuneus Cortex |
| 33 | 3.87 | 32 | -70 | 48 | R | Lateral Occipital Cortex, superior division |
| 31 | 4.3 | -18 | 0 | 26 | L | Left Cerebral White Matter |
